# PASTRI: Resolving Stage-Specific Cell-State Dynamics from Annotated Cell Lineage Trees

**DOI:** 10.64898/2026.08.16.745065

**Authors:** Wenjing Yang, Zizhang Li, Xiaochen Yu, Peng Wu, Xiaoyu Zhang, Chenze Ren, Kehui Liu, Jingyu Chen, Feng Chen, Xionglei He, Jiajun Zhang, Xiaoshu Chen, Jian-Rong Yang

## Abstract

Cellular transitions between phenotypic states are fundamental to development and disease, yet quantitative analysis of their dynamics remains challenging. Here we present PASTRI (*P*hylogenetic *A*djacency-based *S*tate *T*ransition *R*ate *I*nference), a computational framework that infers transition rates from cell lineages/phylogenies annotated with terminal phenotypic states, such as single-cell transcriptomes. We validate PASTRI using simulated lineages and the Caenorhabditis elegans embryonic lineage. Importantly, by leveraging cell pairs at varying phylogenetic distances, PASTRI accurately resolves stage-specific transition rates, circumventing the issue of developmental changes in dynamics. Applied to three cell phylogeny datasets from our lineage-tracing experiments spanning diverse developmental/disease models and tracing systems, PASTRI uncovers rate-limiting steps in the activation of hepatic stellate cells and the differentiation of primordial lung progenitors, as well as attractor states that support cancer cell proliferation. PASTRI thus opens up a venue for dissecting cell state transition dynamics from annotated cell lineage/phylogeny.

## Introduction

The capability of a cell to transition among phenotypically distinct states^1,2^ is a ubiquitous phenomenon in multicellular organisms, governing key biological processes including development, regeneration and disease. For example, in mammalian development, pluripotent cells undergo state transitions marked by changes in patterns of gene expression and developmental potential during lineage fate determination^3–5^. In regenerative contexts, hepatic stellate cells shift from a quiescent to an active state following liver injury, where the activated cells release growth factors that inhibit cell death and stimulate regeneration^6,7^. Many pathologies, from developmental disorders to cancers, are associated with aberrant transitions in cell state. These include the MET(mesenchymal-epithelial transition)-EMT(epithelial-mesenchymal transition) interplay, whose dysregulation leads to congenital defects and increased susceptibility to malignancies and degenerative diseases in adults^8–10^. Quantifying these transition dynamics is therefore essential.

Yet doing so remains challenging. Convential practice has labeled cells or subclones, measured their cellular state (or the state of their descendant cells) at some later time point, and inferred state transition rates from the resulting distribution of labeled cells^11–13^. These efforts struggle with two main obstacles: (i) limited temporal resolution, since labeling and measurement are difficult to repeat at short intervals, and (ii) limited state space granularity, since conventional single-cell measurements capture only a few markers at a time^12^. Recent advances in lineage tracing and single-cell high-throughput assays (especially scRNA-seq) have begun to address these limitations, opening the door to higher-resolution inference of state transition rates. Here, we use ‘cell phylogeny’ to refer to the history of cell divisions revealed by lineage tracing, distinguishing it from ‘cell lineage’, which sometimes denotes a differentiation hierarchy.

The rapid accumulation of cell phylogeny data have motivated several computational methods, including KCA (kin correlation analysis)^1^, CoSpar (coherent, sparse optimization)^14^, Fitch^15,16^ and PhyloVelo (phylogenetic velocity)^17^, which leverage cell phylogenies and scRNA-seq to infer state transition dynamics in the form of cell fate biases, transition matrices, or velocity fields. Despite their promise, these methods share two key limitations. First, the methods assume stable transition rates across the process. This assumption need not hold, however, in two ways — unobserved states may exist, and transition dynamics may change across the course of the biological process under investigation (**Figure 1**). Second, these methods have limited flexibility to incorporate prior knowledge regarding the directionality or rate of state transitions. These two limitations leave the inference of time-varying transition dynamics from cell phylogenies unresolved.

**Figure 1.**
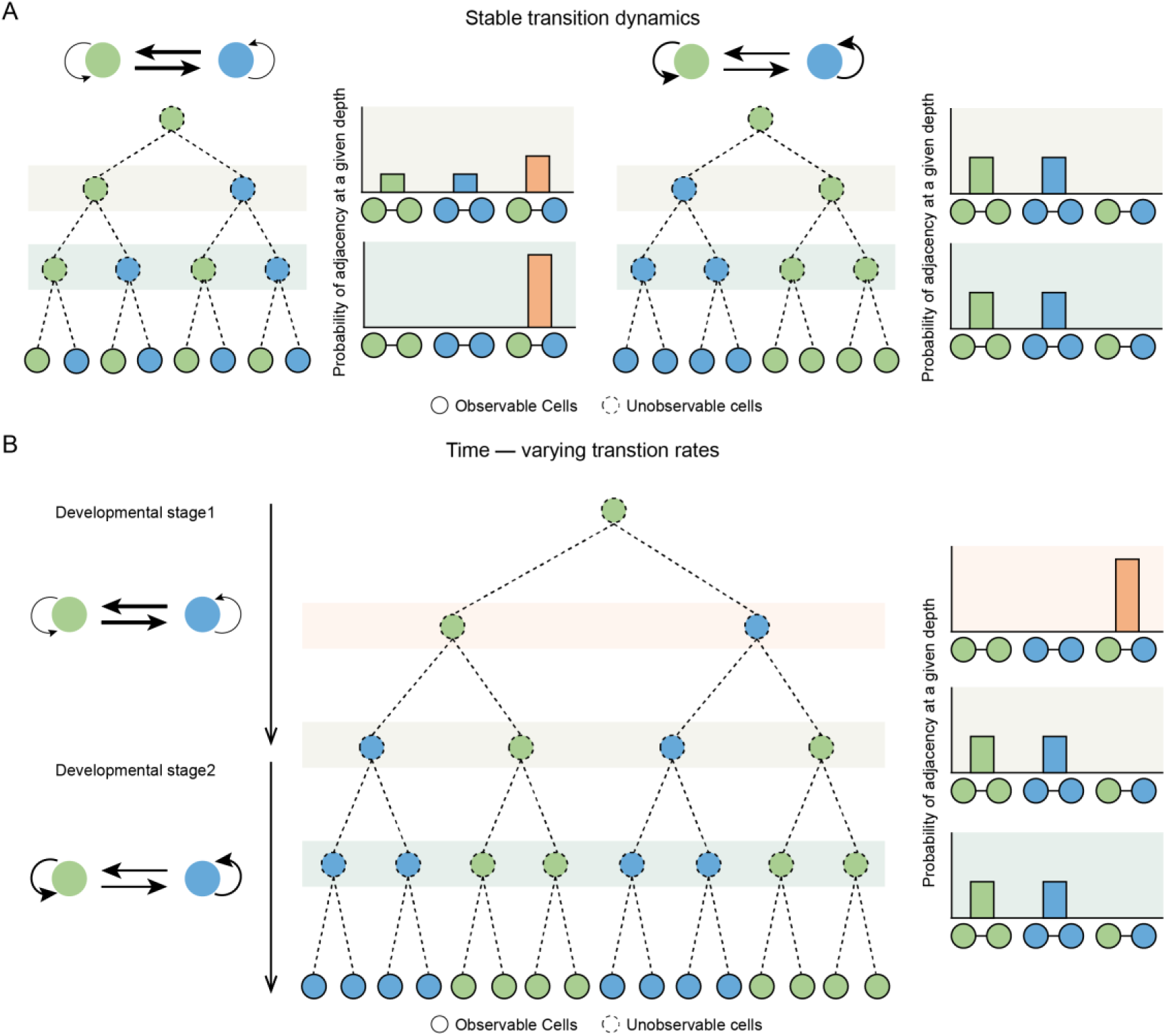
Principles of cell state transition dynamic inference by PASTRI. **(A)** Transition dynamics shape state clustering on the cell phylogeny. The left (right panel shows a two-state model with high transition rate and the expected distribution of terminal cells. The right panel similarly shows a model with low transition rate. Transition dynamics determine the probability of state adjacencies for terminal cell pairs whose MRCAs sits at various depths. **(B)** A two-stage developmental model in which the two states transition at high rate in the early stage 1, but at low rate in the later stage 2. State adjacencies differ between terminal cell pairs whose MRCA are early internal nodes (top right bar plot) and those whose MRCA are later internal nodes (the two bottom bar plots).

To address these two limitations, we developed Phylogenetic Adjacency-based State Transition Rate Inference, or PASTRI. PASTRI infers time-varying transition rates from cell phylogenies, incorporates prior knowledge as inference constraints, and selectively includes phylogenetic adjacencies at various depths on the phylogeny. Applied to simulated and *C. elegans* cell phylogenies with known ground truth, PASTRI infers transition rates more accurately than methods that assume stable rates. We further applied PASTRI to the developmental cell phylogeny of an 8-day *in vitro* directed differentiation from hESCs to hepatic stellate cells (HSC), resolving non-uniform transition rates among HSC activation states. PASTRI generalizes to cell phylogenies of other biological processes reconstructed by various DNA barcode-based methods^18^, including hESC-to-lung-progenitor differentiation and clonal expansion of cancer cell line A549. Together, these results establish cell state transition rates as a plastic feature of living cells and demonstrate that PASTRI can resolve this plasticity from cell phylogenies.

## Results

### Inference of Transition Rates among Phenotypic States using Adjacency on Cell Phylogeny

We infer the cell state transition rates based on cell phylogeny whose terminal nodes (tips) are annotated by cell states (**Figure 1**, **Figure S1** and **Methods**). PASTRI operates on the principle that, given a certain depth, if two cell types are more often adjacent, they should have a high transition rate between them (**Figure 1A**). A pair of terminal cells is considered adjacent at a certain depth *d* if their most recent common ancestor (MRCA) appears at that depth. PASTRI first quantifies the frequency of each state pair for cell pairs adjacent at depth *d*, producing adjacency matrices *A*(*d*). It then infers the transition rate matrix *T* by minimizing the Euclidean distance between adjacencies predicted by a candidate *T* and the observed adjacencies, optionally constrained by prior knowledge of the transition rates (**Figure S1**). PASTRI applies this principle to *A*(*d*) at various depths, leveraging the fact that shallower adjacencies reflect more recent transitions and deeper adjacencies reflect earlier transitions (**Figure 1B**). Algorithmic details of PASTRI are provided.

### Testing PASTRI with Simulated Cell Phylogenies

We first tested PASTRI using simulated cell phylogenies generated from chain-like or branched transition models with stable-reversible dynamics (**Figure 2A-F**). Considering a chain-like transition model, 500 one-division cell phylogenies, each recording one mother cell dividing into two daughter cells, were generated following a predefined matrix of transition rates among five cell states (**Figure 2A**). PASTRI-inferred transition rates based on these simulated phylogenies are correlated with the true (predefined) transition dynamics (Pearson’s R = 0.91, *P* < 10⁻⁴; **Figure 2B**). The inferred transition rates recapitulated the truth with a Euclidean distance of 0.26, whereas inference based on 500 randomized phylogenies created by shuffling the terminal cell state labels yielded a mean Euclidean distance of 0.42 with the truth (*P* < 0.04, Permutation test; **Figure 2C**). Comparable results were obtained with branched transition models (**Figure 2D-F**) and with stable, irreversible dynamics (**Figure S2**). These results confirm PASTRI’s accuracy.

**Figure 2.**
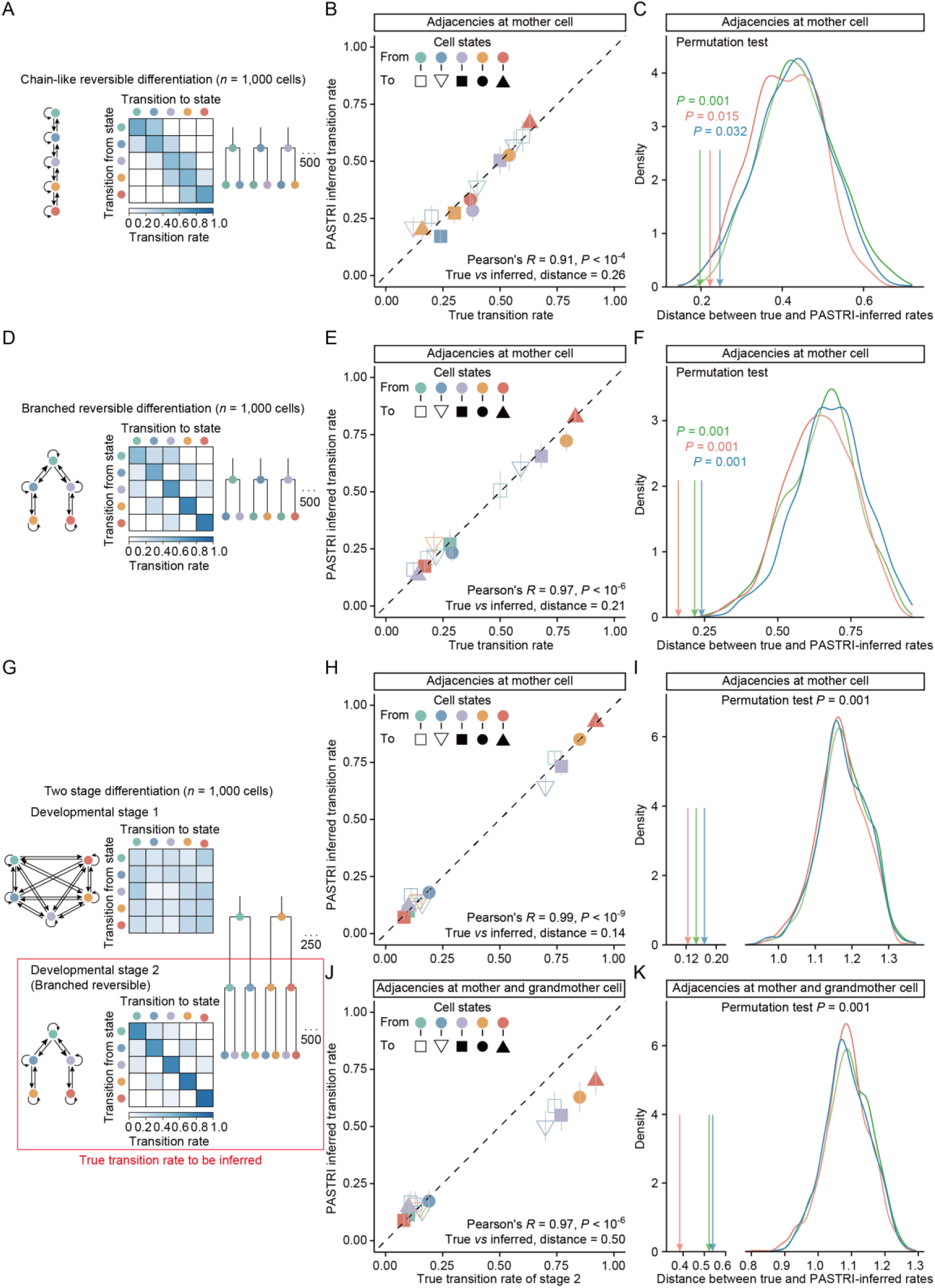
Testing PASTRI with simulated cell phylogenies. **(A)** A stable-reversible chain-like transition model, depicted as a network or matrix, was used to simulate 500 phylogenies of 2 terminal cells each. **(B)** A typical result from one simulation. The true (*x*) and inferred (*y*) transition rates from state *i* (point color) to state *j* (point symbol) are compared. The bottom right corner shows the Pearson’s correlation test result and Euclidean distance between the true and inferred transition rates. **(C)** A colored arrow marks the observed Euclidean distance between the true and PASTRI-inferred transition rates. A curve of the same color shows the expected probability density of the Euclidean distance over 1,000 sets of phylogenies (500 per set) obtained by shuffling the terminal cell states. Three pairs of observed and expected Euclidean distances, from three independent simulation rounds, are shown in three colors. (**D-F**) Same as A-C, except that they are based on a stable-reversible branched transition model. (**G**) A two-stage transition model with a branched reversible stage 2 was used to simulate 250 phylogenies, each undergoing two rounds of cell divisions and producing 4 terminal cells. **(H-I)** Same as B-C, except that PASTRI used mother-level adjacencies only, and was compared to the stage-2 truth. **(J-K)** Similar to H-I, except that the inference was made using adjacencies at mother and grandmother cells.

To assess how time-varying transition dynamics affect transition rate inference, we conducted another test of PASTRI’s performance on a two-stage differentiation model. Using one distinct predefined transition rate matrix for each round of division, we simulated 250 cell phylogenies, each recording two rounds of cell division, from 250 grandmother cells, to 500 mother cells, and then 1,000 terminal cells (**Figure 2G**). PASTRI inferred the stage-2 (second division) transition matrix from mother-level adjacencies (Euclidean distance 0.14, *P* < 0.001, Permutation test; **Figure 2H-I**) or from both mother and grandmother-level adjacencies (Euclidean distance = 0.50, *P* < 0.001, Permutation test; **Figure 2J-K**). When adjacencies at both mother and grandmother cells were combined, the inferred rates resemble neither stage’s true matrix, with a Euclidean distance of 0.50 from the stage-2’s truth and worse still for the stage-1’s (**Figure S3A-B**). This deterioration holds across the transition model types tested (**Figure S3C-H**).

To assess how cell subsampling, due to failure to capture cells or barcodes, interferes with PASTRI, we simulated ten phylogenies each with 50,000 terminal cells using a random bifurcation model (see **Methods**). In the simulation, two distinct rate matrices govern the cell state transitions at the early and late stages (**Figure S4A**). We randomly selected 50%, 20%, or 10% of the tips (25000, 10000, or 5000 cells) to construct a partial phylogeny, recalculating the internal cells’ depths from the partial phylogeny to mimic cell subsampling. We compared the PASTRI results from the full and partial phylogenies with the true transition rates, using Euclidean distances. As expected, cell subsampling reduces inference accuracy (**Figure S4B**). Yet PASTRI outperforms algorithms that assume stable rates (**Figure S4B**). This advantage follows from PASTRI’s selective inclusion of adjacencies at appropriate depths to infer time-varying transition rates, yielding accurate inference of the corresponding developmental stage. We hereinafter focus on transition dynamics near the terminal cells, where PASTRI’s ability to selectively include adjacencies at shallower depths enables the inference of recent transitions, the most informative target for PASTRI.

### PASTRI Accurately Infer Cell State Dynamics in Cell Phylogenies of *C. elegans*

We benchmark PASTRI on cellular state transitions defined by the expression of functional markers in real cell phylogenies of *C. elegans*^19^. From the Expression Pattern In Caenorhabditis (EPIC) database^20,21^, 213 cell phylogenies (>200 tips) each annotated with spatial-temporal expression pattern of one of 115 marker genes, some with biological replicates, were extracted (**Figure 3A**). An example annotated phylogeny for the gene *sma-9* encoding a transcription factor with dual coactivator and corepressor activities is shown (**Figure 3B**). On these phylogenies, a cell’s depth is calculated based on the number of additional cell divisions recorded for its descendants, normalized to the range 0 (tip) to 1 (root) (*y* axis in **Figure 3B**; See **Methods**).

**Figure 3.**
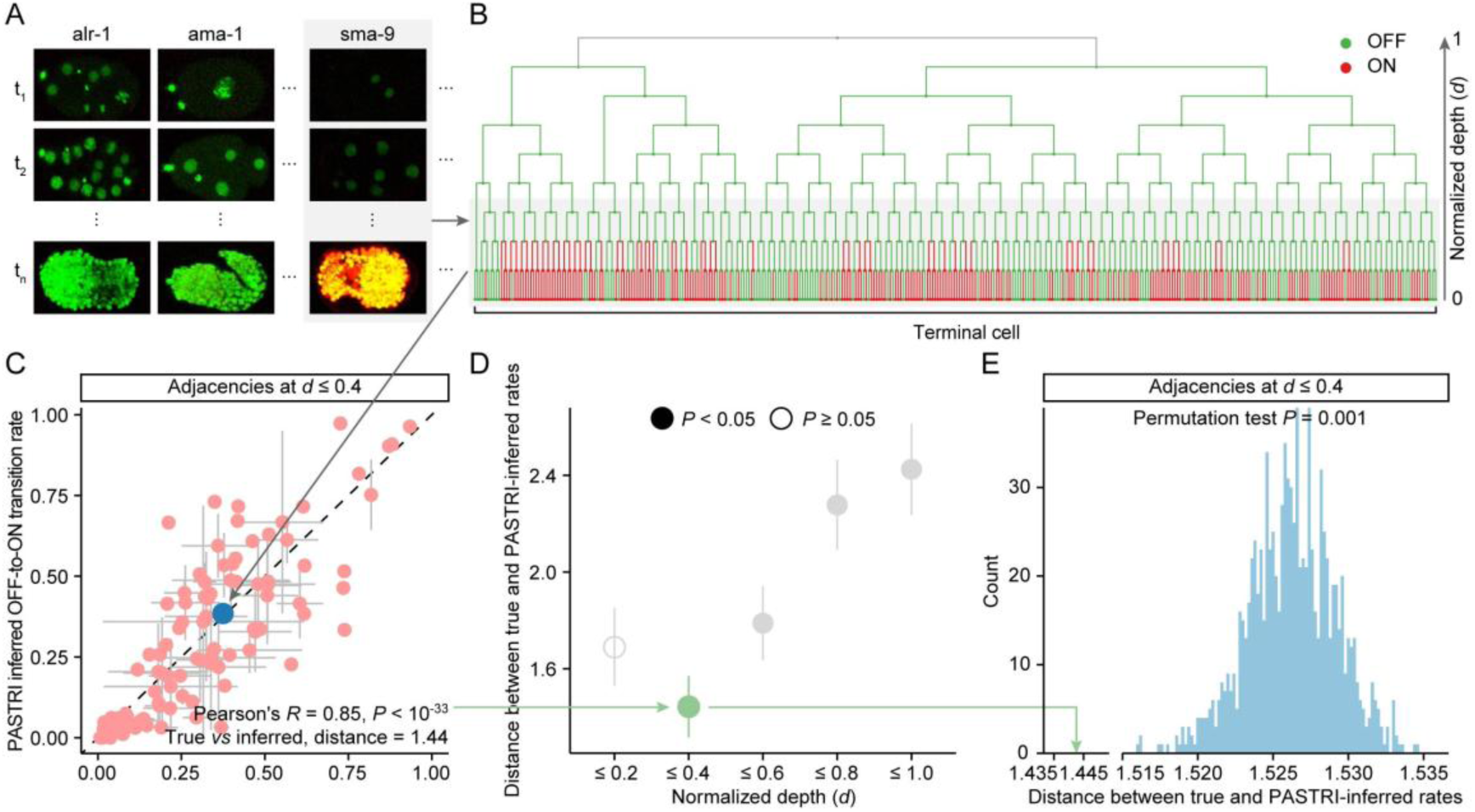
Evaluating PASTRI with cell phylogenies of *C. elegans*. **(A)** The EPIC database^20^ provides time-lapse movies that trace the cell lineage-specific expression patterns of marker genes during *C. elegans* development. **(B)** We extracted the expression status (ON or OFF) of each marker gene together with the traced cell phylogeny (see **Methods**). The example shows the gene *sma-9* on a cell phylogeny traced to 353 terminal cells. **(C)** Using adjacencies up to *d* ≤ 0.4, PASTRI inferred OFF-to-ON transition rates (*y*) are compared with the true transition rates (*x*) (see **Methods**). The blue dot indicates the example gene *sma-9*. Gray lines represent the standard error between biological replicates. Pearson’s correlation coefficient and *P* value appear at the bottom, together with the Euclidean distance between the inferred and true rates. **(D)** The Euclidean distance between the inferred and true rates (*y*) is shown for PASTRI inference across adjacencies up to various *d* (*x* axis). The vertical lines represent standard deviation by bootstrapping the genes 1,000 times. Solid (*P* < 0.05) or hollow (*P* ≥ 0.05) circles indicate the statistical significance (Permutation tests). See also **Figure S5**. **(E)** The arrow marks the observed Euclidean distance between the inferred and true rates. The blue histogram shows the expected probability density of the Euclidean distance over 1,000 phylogenies obtained by shuffling the terminal cell states. *P* values from permutation tests are indicated on top

For each gene, the true transition rate from inactive (OFF) to active (ON) state was calculated by tallying mother-daughter state changes in the real cell phylogeny up to various depths (**Figure 3C**. See **Methods**). PASTRI inferred the same transition rates, using the cell phylogenies and the terminal cell (but not internal cell) annotations of ON/OFF states of these 115 genes, by considering adjacency matrices up to various depths (**Figure 3C-D** and **Figure S5A-B**. See **Methods**). Averaging results from replicated phylogenies of the same gene, we obtained transition rate matrix for cellular states defined by each of the 115 genes. We quantified PASTRI’s accuracy as the Euclidean distance between the inferred and the true OFF-to-ON transition rates (**Figure 3C** and **D**), and assessed statistical significance by comparing the inference from the true cell phylogeny to that from randomized phylogenies with shuffled terminal cell state labels (**Figure 3E** and **Figure S5B**. See **Methods**).

We found that limiting PASTRI to proximal adjacencies improved its accuracy and statistical significance until the sparsity of the remaining adjacencies hindered better inferences (**Figure 3D**). A closer look at the most accurate results (**Figure 3C**, Euclidean distance = 1.44) revealed that PASTRI inference is reasonably good regardless the values of the true transition rate, and outperforms other algorithms that assume stable transition rates (**Figure S5C**). For example, *sma-9* exhibits a true OFF-to-ON transition rate (scaled to a time unit of *d*=1) of 0.375 during the last three rounds of divisions (*d* ≤ 0.4) recorded. Using the cell phylogeny and only the terminal states, PASTRI gives an accurate inference of 0.385 (**Figure 3C**, the blue dot). On the contrary, when the more distal adjacencies were used in PASTRI, an increasingly stronger overestimation of the transition rates was observed (**Figure S5A-B**). This is likely caused by the slower cell state transition during the early development of *C. elegans*, a phenomenon that cannot be inferred from the tips of the cell phylogeny. Collectively, these results supported PASTRI’s capacity and highlighted that stage-specific inference of transition rates is more informative than overall inference in real developmental cell phylogenies.

### PASTRI Identifies a Rate-limiting Transition during Early Activation of Hepatic Stellate Cells

Hepatic stellate cells (HSCs) derive from human embryonic stem cells (hESCs) through a multi-step differentiation program, and their activation drives liver fibrogenesis^22,23^. We applied PASTRI to cell phylogenies reconstructed by the substitution mutation-aided lineage-tracing (SMALT) system^24^ to identify the rate-limiting transition in this program. To this end, we constructed a lineage tracer hESC cell line consisting of the SMALT system with a transcribed lineage barcode, enabling simultaneous cell-phylogeny and cell-state readout via single-cell RNA-seq (**Figure 4A**. See **Methods**). This cell line was constructed with three major steps (**Figure S6A**): (i) genomic integration of a SMALT inducible mutator; (ii) single-copy genomic integration of the transcribing lineage barcode targeted by the SMALT mutator; and (iii) optimization of the stability and efficiency of the SMALT system in hESC (see **Methods**, **Figure S6B-G**).

**Figure 4.**
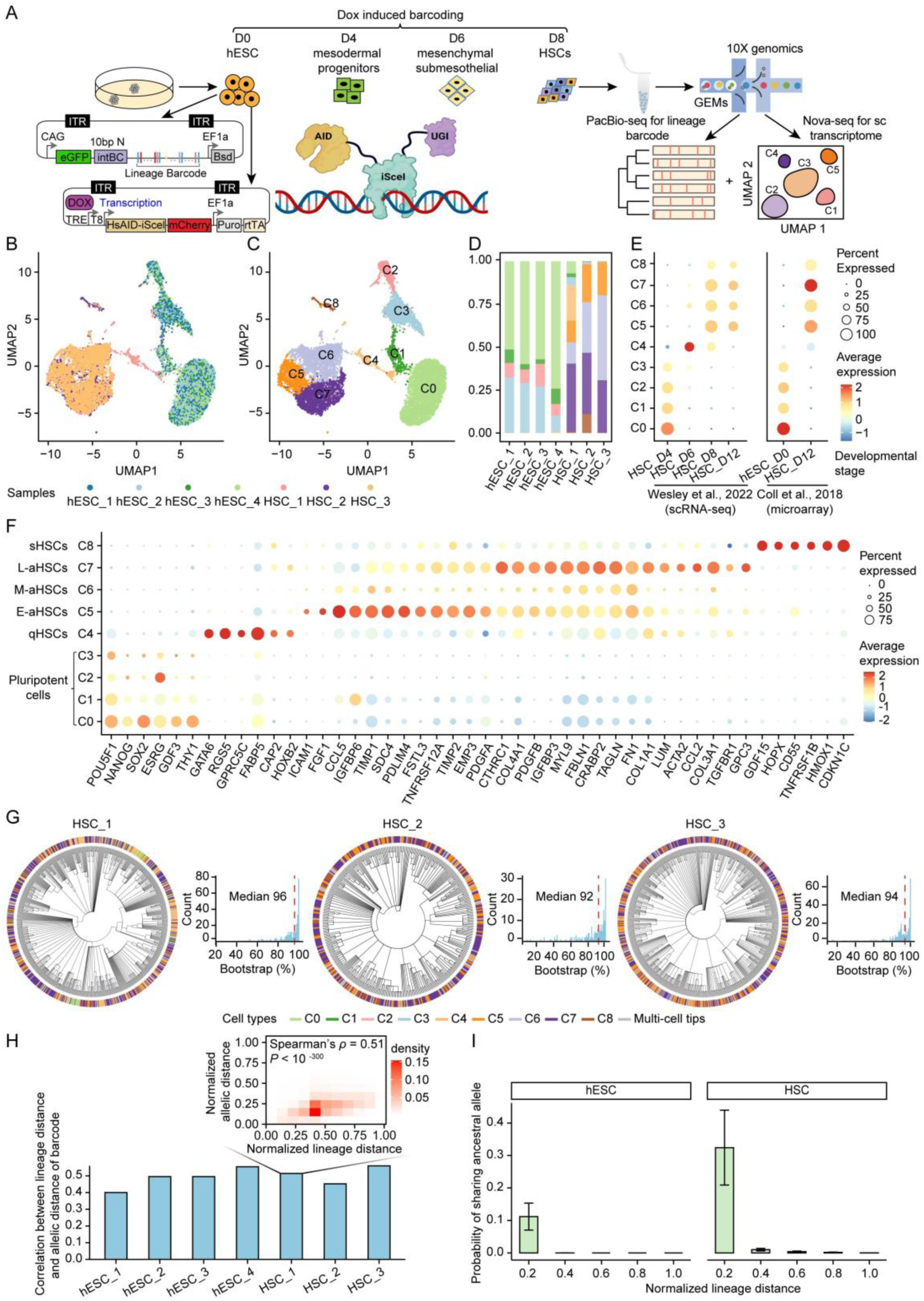
*In vitro* hESC-to-HSC differentiation reconstructed from single-cell transcriptomes and SMALT lineage barcodes. **(A)** Schematic of the experimental workflow. Directed differentiation from lineage tracer hESCs to HSCs was carried out for eight days, along with simultaneous lineage tracing by the SMALT system. The resulting colonies were assayed for single-cell transcriptomes by Nova-seq and lineage barcodes by PacBio HiFi-seq, which were used to reconstruct cell phylogenies with single-cell transcriptomes assigned to tips. **(B)** UMAP embedding of 33,438 single-cell transcriptomes, colored by samples. **(C)** Nine subclusters (cell states) resolved by clustering of the single-cell transcriptomes. **(D)** Percentage of cells in each cell state (*y* axis) across the seven samples (*x* axis; 3 differentiating and 4 non-differentiating). Cell states colored as in **C**. **(E)** Average expression (color) and percentage of expressed cells (dot size) for differentially expressed gene sets of each developmental stage (*x* axis; defined by either single-cell transcriptomics or microarray) across the nine cell states (*y* axis). **(F)** Average expression (color) and percentage of expressed cells (dot size) for marker genes of pluripotent cells and HSCs (*x* axis) across the nine cell states (*y* axis). Pluripotent cells (C0/C1/C2/C3) express *POU5F1*/*OCT4* and other pluripotency-associated genes; quiescent HSCs (C4, qHSCs) express *GATA6* and *RGS5*^51,52^; activated HSCs (C5/C6/C7) express *CCL5* (early activated marker), *TIMP1*, *FN1*, and *COL3A1* (late activated marker)^32,52–54^; senescent HSCs (C8) express GDF15, HOPX, CDKN1C, and HMOX1^28,55–57^. **(G)** Cell phylogenies of the three differentiating samples, with terminal tips colored by cell state. Inset histograms show the bootstrap support distribution across each phylogeny’s internal nodes. See also **Figure S8D** for non-differentiating samples. **(H)** Correlation between normalized lineage distance (depth of MRCA of two cells, divided by sample’s maximal depth) and normalized barcode allelic distance (target sites differing from each other, divided by the maximum). Inset: two-dimensional density plot for one sample. **(I)** Probability of shared ancestral alleles between two single-cell tips (*y* axis) decreases with normalized lineage distance (*x* axis; defined as in **H**). Error bars indicate standard error across samples.

Using this cell line, we reconstructed the cell phylogeny of human embryonic stem cells (hESCs) undergoing *in vitro* directed differentiation into HSCs. We selected this model for three reasons. First, *in vitro* culture enabled control of cell counts during differentiation, capturing 10.8–30.5% of terminal cells for sequencing, exceeding the ≤1% capture rate of *in vivo* studies^24,25^. Second, this *in vitro* model closely mimics *in vivo* HSC developmen^22,23^, with ∼60% of cells reaching an HSC fate after 8 days of directed differentiation (**Figure S7A–D**). Third, *in vitro* culture co-initiated differentiation and SMALT mutator expression, so the lineage barcode began recording at the onset of differentiation. Parallel non-differentiating cultures maintaining their hESC states are used as controls (see **Methods**).

We sampled three differentiating and four non-differentiating cultures exhibiting normal morphology and uniformly strong mCherry fluorescence (**Figure S8A**), each subjected to simultaneous PacBio sequencing of the lineage barcode and NovaSeq scRNA-seq (**Figure 4A**. See **Methods**). We obtained 33,438 high-quality single-cell transcriptomes. Differentiating and non-differentiating samples differed clearly in their transcriptomes (**Figure 4B**). Clustering resolved the cells into nine subclusters (**Figure 4C**), which showed distinct distributions between the two conditions (**Figure 4D**) and matched known cellular states of directed differentiation^22,26^ (**Figure 4E**). Marker genes for pluripotent cells and HSC development assigned the nine subclusters to four groups: pluripotent cells (C0-C3), quiescent HSCs (C4, qHSCs), activated HSCs (C5-C7), and senescent HSCs (C8) (**Figure 4F**; see caption for marker details). Mapping the four groups onto the expected differentiation trajectory confirmed the *in vitro* differentiation protocol.

Targeted PacBio HiFi sequencing identified accumulated mutations in the lineage barcodes of 8,474 cells (see **Methods**). Over 86% of the mutations were CG-to-TA, consistent with the cytidine deaminase activity of the SMALT mutator (**Figure S8B**). Mutations accumulated at 4.73–10.78 per cell across samples (**Figure S8C**). These lineage barcodes reconstructed maximum-likelihood cell phylogenies of 451–2,203 single cells per sample (**Figures 4G** and **S8D**). The phylogenies had median bootstrap support >92% across their internal nodes (**Figures 4G** and **S8D**), and their lineage/phylogenetic distance correlated with barcode allelic distance (**Figures 4H** and **S8E**). Closely related cells shared ancestral barcode alleles (**Figure 4I**) and showed transcriptomic similarity (**Figure S8F**). These evidence validated the phylogenies as reliable substrates for PASTRI inference of HSC differentiation.

We applied PASTRI to the differentiating phylogenies to infer transition rates between HSCs captured on day 8, assuming irreversible transitions between early (E-aHSCs), intermediate (M-aHSCs), late aHSCs (L-aHSCs) and senescent HSCs (sHSCs)^27,28^. PASTRI inferences at adjacencies of normalized depth *d* ≤ 0.5 showed the smallest Euclidean distance between transition matrices across biological repeats and the highest statistical significance (**Figures 5A-B** and **S9A**), and outperformed other algorithms that assume stable transition rate (**Figure S9B**). Transition rates at this depth spanned two orders of magnitude across HSC state pairs, including a transition rate of 0.04 from E-aHSCs to M-aHSCs, 0.84 from M-aHSCs to L-aHSCs, and 0.60 from E-aHSCs to L-aHSCs (**Figure 5C**). The E-aHSC to M-aHSC transition was the rate-limiting transition in HSC activation: its rate (0.04) was more than 20-fold lower than the M-aHSC to L-aHSC rate (0.84). E-aHSCs respond to liver injury signals without fully committing to a fibrogenic phenotype^29,30^. This slow transition aligns with prior reports that E-aHSCs must undergo metabolic and phenotypic changes before transitioning to the proliferative, fibrogenic M-aHSC state^31^. At M-aHSC, aHSCs acquire proliferative and fibrogenic activity that supports the rapid L-aHSC transition^27,32–34^. Rapid transitions coincided with smaller transcriptomic divergence (**Figure 5D**) and smaller DEG counts (**Figure 5E**), consistent with the biological interpretation of PASTRI’s transition rates. Together, these results demonstrated the power of PASTRI in revealing intricate cell state transition dynamics.

**Figure 5.**
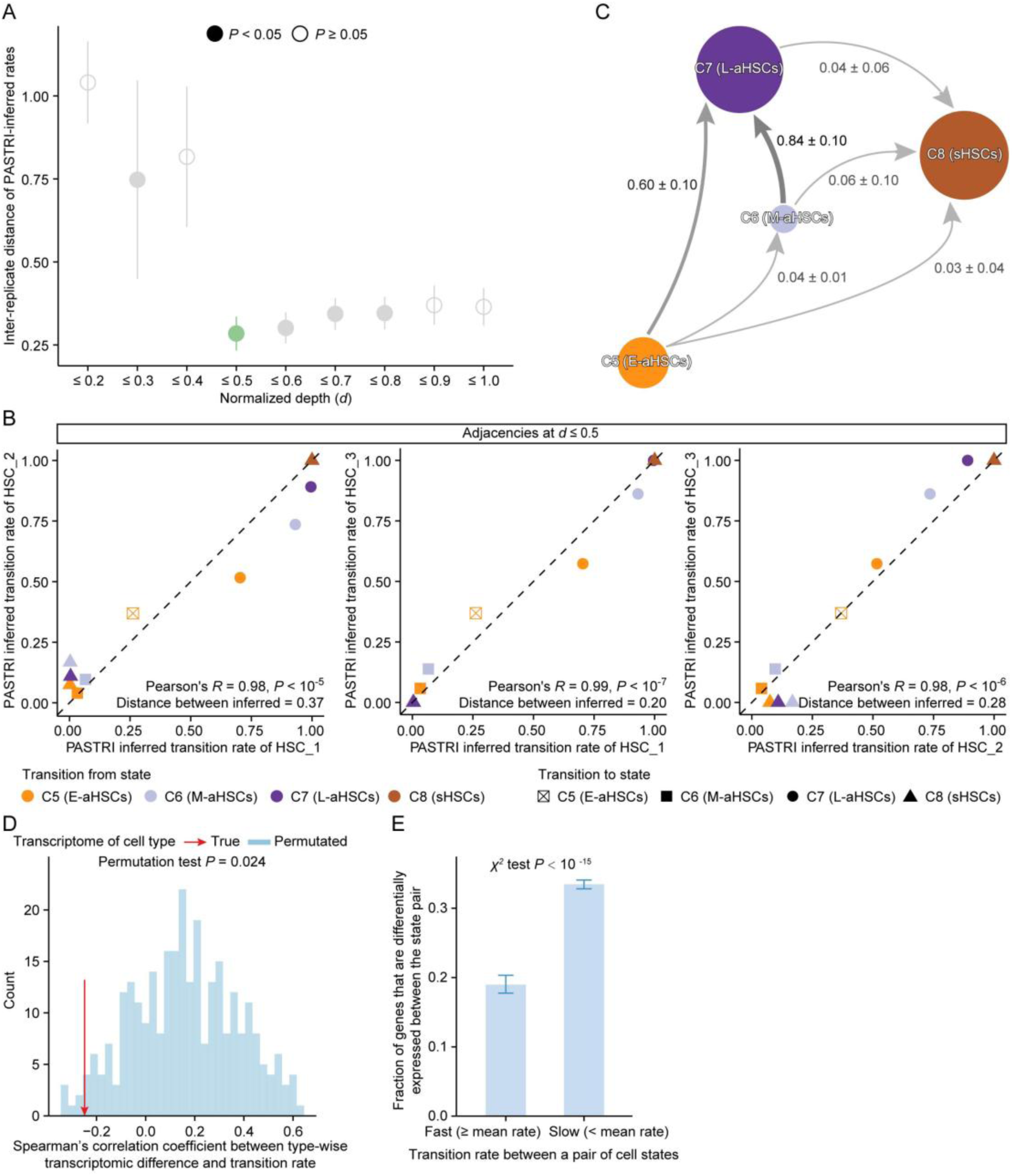
PASTRI infers HSC activation transitions from SMALT-reconstructed phylogenies. **(A)** Among-replicate reproducibility (*y* axis) of PASTRI inferences when using adjacencies up to various *d* (*x* axis). Vertical lines indicate standard errors between biological replicate pairs; solid and hollow circles mark permutation *P* values < 0.05 and ≥ 0.05, respectively. See also **Figure S9A**. **(B)** Pairwise reproducibility between biological replicates, based on PASTRI inference using adjacencies up to *d* ≤ 0.5. Start and end states distinguished by color and shape. Pearson’s correlation coefficient, *P* value, and Euclidean distance are indicated for each pair. **(C)** Cell state transition graph averaged from the three biological replicates, inferred by PASTRI using adjacency matrices up to *d* ≤ 0.5. **(D)** Negative correlation between PASTRI-inferred transition rates and transcriptomic divergence between cell state pairs (red arrow), compared with the random expectation gauged by 287 (= 4! × 3! × 2! − 1) randomized controls (histogram). Controls are created by shuffling transition rates within each starting state. **(E)** Cell state pairs with fast transition (left) carry fewer differentially expressed genes than those with slow transitions (right). Error bars indicate standard deviation from 1,000 bootstraps of the genes. The *P* value from a *χ*^2^ test is indicated on top.

### PASTRI Elucidates State Transition Dynamics during hESC Differentiation to Lung Progenitor Cells

We applied PASTRI to CRISPR-Cas9-based cell phylogenies reconstructed for an *in vitro* directed-differentiation model of hESCs into primordial lung progenitors (PLP)^35^. Twelve major functional clusters were identified within the cells of three independent differentiating cultures^35^, which belonged to three cell fate commitment stages: pluripotent cells (C1:NANOG^hi^POU5F1^hi^, C2:NANOG^low^POU5F1^hi^, C3:NANOG^low^POU5F1^low^, C4:NANOG^hi/low^POU5F1^hi^), lung progenitor cells (C5: CD47^hi^, C6:CD47^low^, C7:GATA6^hi^SHH^hi^CD47^low^, C8:GATA6^low^NKX2-1^neg^SHH^neg^CD47^neg^, C9:GATA6^hi^NKX2-1^hi^CD47^hi^, C10:GATA6^hi^), and spontaneous cells (R1, R2). The corresponding cell phylogenies have a depth of ∼13 and contained thousands of terminal cells with mixed cell states in hierarchical subclones (an example in **Figure 6A**). Assuming irreversible transitions among the progenitor cell states^35^, we applied PASTRI to the three hESC-PLP phylogenies and inferred transition rates among the six lung progenitor cell states (see **Methods**). We used adjacencies at *d* ≤ 0.6, which gave the smallest average Euclidean distance between transition matrices across biological repeats and the highest statistical significance against random expectations (**Figures 6B** and **S9C**). At this depth, PASTRI-inferred transition rates were reproducible across samples (**Figure 6C**) and heterogeneous across cell states (**Figure 6D**); PASTRI also outperformed other algorithms that assume stable transition rate (**Figure S9D**). PASTRI inferred a transition rate of 0.01 from C7 to C8, 0.35 and 0.23 from C8 to C9 and C10, and 0.16 and 0.19 from C7 to C9 and C10 (**Figure 6D**). The slow C7 to C8 transition identifies C8 as the rate-limiting intermediate for the formation of PLP (C9, marked by NKX2-1^hi^) or other endoderm-derived tissues (C10)^36^ from an endoderm stage (C7). This finding aligns with the temporal accumulation of fibroblast growth factors (FGF) past a critical threshold for specific pulmonary cell fate^36^, supporting PASTRI’s reliability across a different lineage tracing system and biological contexts.

**Figure 6.**
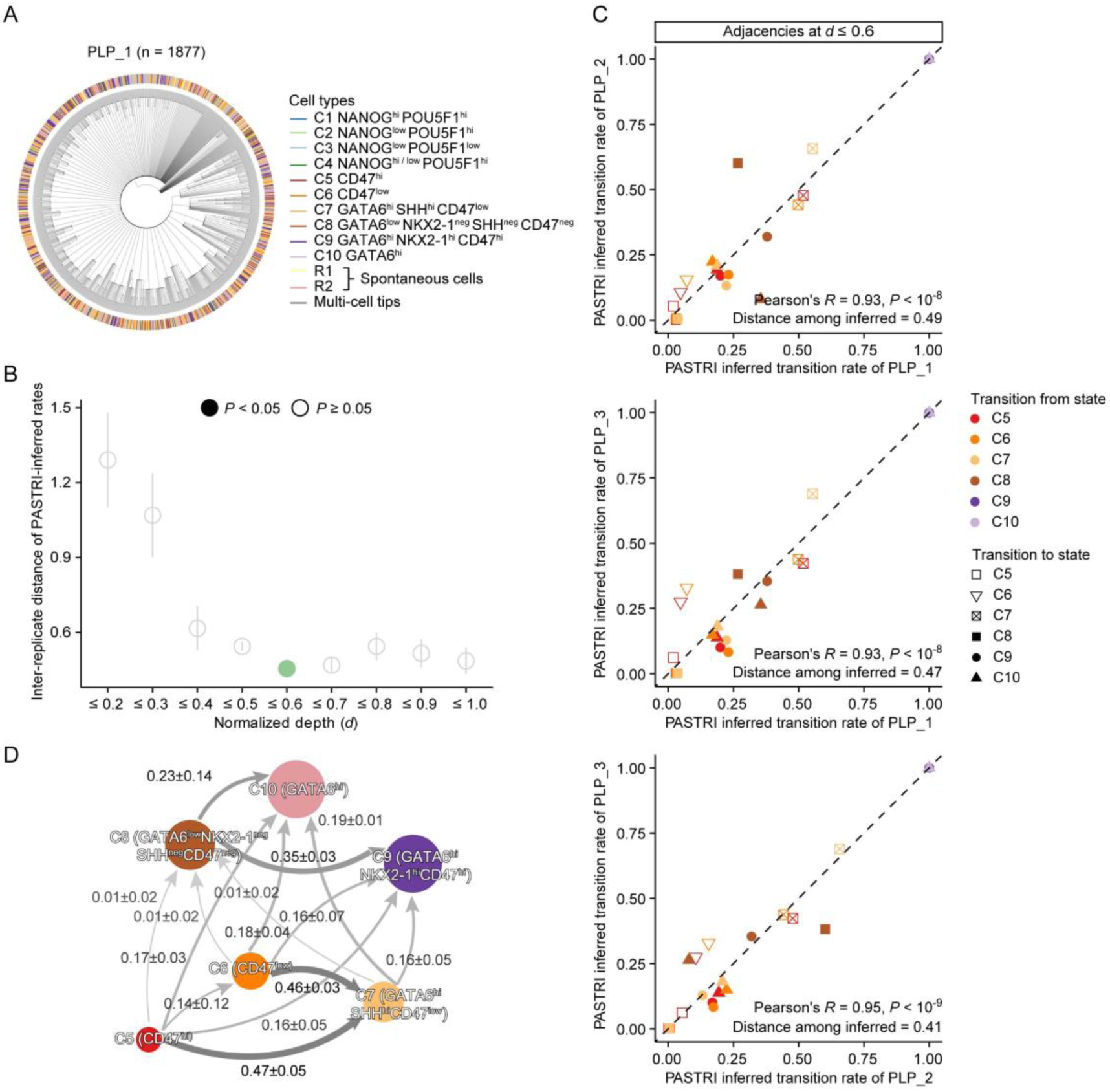
PASTRI infers lung progenitor cell state transitions from scGESTALT-reconstructed hESC-PLP phylogenies. **(A)** Annotated cell phylogeny of an hESC-PLP directed differentiation (1,877 single cells)^35^. **(B)** Among-replicate reproducibility (*y* axis) of PASTRI inferences versus adjacency depth *d* (*x* axis). Vertical lines indicate standard errors between biological replicate pairs; solid and hollow circles mark permutation *P* values < 0.05 and ≥ 0.05, respectively. See also **Figure S9C**. **(C)** PASTRI-inferred transition rates at *d* ≤ 0.6, plotted for pairwise reproducibility between biological replicates. Start and end states distinguished by color and shape. Pearson’s correlation coefficient, *P* value, and Euclidean distance are indicated for each pair. **(D)** Cell state transition graph averaged from the three biological replicates, inferred by PASTRI at *d* ≤ 0.6.

### PASTRI Reveals Attractor States Supporting Proliferation of Cancerous Cells

Beyond differentiation during development, cell state dynamics during cancer somatic evolution carry high biomedical importance — driving tumor progression, drug resistance, and invasive potential. We applied PASTRI to SMALT-based cell phylogenies obtained from *in vitro* cultures of human lung cancer cells A549. Three samples totaling 17,327 high-quality single-cell transcriptomes clustered into six distinct cell states (**Figure 7A–D**): a proliferative core of C0 (chromatin assembly), C1 (cell-cycle), and C2 (microenvironment sensing), and three terminal states C3 (non-coding RNA-high), C4 (indistinct), and C5 (mitochondrial-high) (see **Figure 7** caption for marker details).

**Figure 7.**
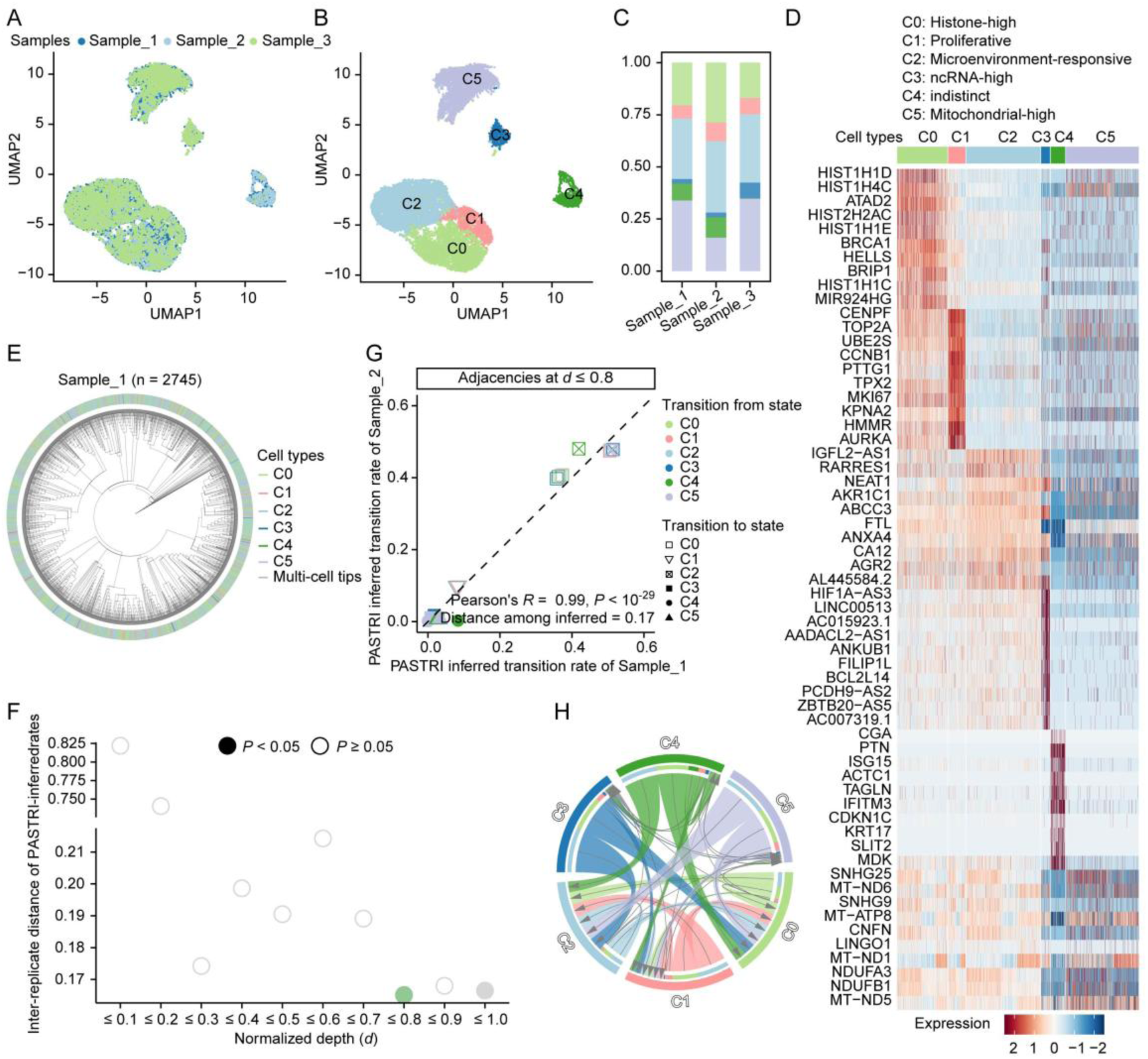
PASTRI infers cancer cell state transitions from SMALT-reconstructed phylogenies during *in vitro* A549 clonal expansion. **(A)** UMAP embedding of the 17,327 single-cell transcriptomes, colored by A549 sample. **(B)** Six subclusters (states) resolved by clustering of the single-cell transcriptomes. **(C)** For each of the three samples (*x* axis), the *y* axis shows the percentage of cells in each state. Cell states colored as in **B**. **(D)** Heatmap of the top-ten most-expressed genes (*y* axis) per cell state across all single cells (columns), with cells grouped by state (labeled above). Marker assignments: C0 (Histone-high) by core histone genes (*HIST1H1D*, *HIST1H4C*, *HIST1H1C*)^58^; C1 (Proliferative) by cell-cycle genes (*CENPF*, *CCNB1*, *TOP2A*)^59,60^; C2 (Microenvironment-responsive) by tumor microenvironment-associated genes (*IGFL2-AS1*, *RARRES1*, *NEAT1*)^61–63^; C3 (Non-coding RNA-high) by ncRNAs involved in tumor microenvironment regulation (*HIF1A-AS3*, *LINC00513*, *AADACL2-AS1*)^64–66^; C5 (Mitochondrial-high) by mitochondrial function genes (*MT-ND6*, *MT-ATP8*, *NDUFA3*)^67–69^; C4 (indistinct) lacked clear marker expression. **(E)** One annotated cell phylogeny of the *in vitro* clonal expansion of A549 cell. **(F)** Among-replicate reproducibility (*y* axis) of PASTRI inferences versus adjacency depth *d* (*x* axis). Solid and hollow circles mark permutation *P* values < 0.05 and ≥ 0.05, respectively. See also **Figure S9E**. **(G)** PASTRI-inferred transition rates at *d* ≤ 0.8, plotted for pairwise reproducibility between Sample_1 (*x* axis) and Sample_2 (*y* axis). Start and end states distinguished by color and shape. Pearson’s correlation coefficient, P value, and Euclidean distance between transition matrices are shown. **(H)** Circos plot of transition rates between cell states (colored on the outer circle). Ribbon width at the originating end marks the transition rate; the arrow marks direction.

The corresponding cell phylogenies had an average depth of 33, intermixing cells of different states (**Figure 7E**). Compared with non-cancer samples (hESC, HSC, and lung progenitor), cancer cells accumulated more barcode mutations, giving the phylogenies higher resolution^37^. We excluded sample 3 from PASTRI analysis because > 20% of the terminal nodes on the phylogeny were not single cells. Adjacencies at *d* ≤ 0.8 gave the smallest Euclidean distance between samples 1 and 2’s transition matrices and the highest statistical significance (**Figure 7F-G** and **Figure S9E**), and also outperforms algorithms that assume stable transition rates (**Figure S9F**). Compared to the developmental phylogenies, the larger depth of the optimal inference indicated slower transitions, while the more promiscuous transition (**Figure 7H**) revealed greater plasticity in cancer cells. The quantitative transition rate map revealed slow transitions to C3, C4, and C5, and identified C0, C1, and C2 as attractor states readily reached from other states (**Figure 7H**). C0, C1, and C2 thus form the core functional programs sustaining tumor survival and expansion — C0 enabling chromatin assembly for DNA replication, C1 driving proliferation, and C2 remodeling the tumor microenvironment. The plasticity of these three states underpins tumor adaptability and invasive potential^38,39^. Extending PASTRI’s reach from development to cancer somatic evolution, this analysis identified attractor states that sustain tumor progression.

## Discussion

Here, we proposed PASTRI as a method to infer cell state transition rates directly from cell phylogenies with phenotypic states annotated at terminal nodes. Through tests on simulated and *C. elegans* cell phylogenies with known ground truth, PASTRI recovered transition rates that matched expected values. Applied to SMALT-reconstructed phylogenies of hESC-to-HSC differentiation, PASTRI identified the rate-limiting transition from early to intermediate activated HSCs. In scGESTALT-based phylogenies of hESC-to-lung-progenitor differentiation, PASTRI identified C8 as a rate-limiting intermediate; the same framework extended to SMALT-based phylogenies of A549 cancer cell clonal expansion to reveal attractor states sustaining tumor progression. These applications establish PASTRI as a quantitative framework for resolving cell state dynamics at single-cell resolution, complementing transcriptomic snapshots with phylogeny-encoded temporal information.

There are three caveats in our study that should be discussed. First, PASTRI’s accuracy depends on cell composition, phylogeny topology, and cell distribution on the phylogeny. Our permutation test controls only the third factor, yielding a conservative significance estimate. Second, PASTRI incorporates prior knowledge of transition dynamics (such as irreversibility). Errors in such assumptions reduce accuracy, motivating functional validation of transitions with low-confidence or poor reproducibility. Third, the PASTRI’s accuracy scales with cell sampling density (see **Figure S4**). PASTRI results nonetheless remains reproducible for phylogenies sampling <0.1% cells^40^ (**Figure S10**). As denser sampling gains recognition^18,35,41–43^, PASTRI would resolve cell fate transitions at increased resolution.

Our hESC-to-HSC dataset comprises three differentiating and four non-differentiating cell phylogenies, with sampling density at 10.8–30.5% of differentiating cells. Combined with the SMALT system’s temporal resolution, these cell phylogenies enable fine-grained cell-fate mapping of hESC-to-HSC differentiation. PASTRI inferences were most accurate at *d* ≤ 0.5, indicating that adequate cell sampling is required for reliable transition rate estimation. If the sampling rate were halved, cousin cells on the original phylogeny would reclassify as sister cells; at extreme under-sampling (one cell per subtree at *d* ∼ 0.5), PASTRI inference reduces to adjacencies at 0.5 < *d* < 0.6 of the original phylogeny, where accuracy likely degrades. Denser cell sampling would enable PASTRI to fully realize its stage-specific inference capability.

Beyond the shown applications, two features distinguish PASTRI from existing tree-based methods. First, PASTRI infers stage-specific transition rates without assuming a single time-invariant rate matrix, an assumption shared by existing methods, thereby enabling quantitative re-examination of conclusions drawn under the fixed-rate assumption. Second, as a platform-agnostic and open-source framework, PASTRI can be retroactively applied to the rapidly growing corpus of cell lineage tracing datasets. We anticipate PASTRI will accelerate the construction of quantitative, time-resolved maps of cell state dynamics, akin to Waddington’s landscape.

## Methods

### The Phylogenetic Adjacency-based State Transition Rate Inference (PASTRI) algorithm

PASTRI analyze a cell phylogeny in the form of a tree (i.e., an acyclic directed graph), in which each node represents one cell, and each branch represents a descendant relationship from an ancestral cell to one of its descendent cells. The cells/nodes can be categorized into internal or terminal based on whether their further divisions were recorded by the phylogeny. Ideally, the internal nodes should be bifurcating, reflecting the division of a mother cell into two daughter cells. However, some multifurcation may occur in real data because lineage tracing experiments may not provide sufficient temporal resolution. All terminal cells are labeled by their functional or transcriptomic state, which is usually the expression level or status of one or more marker genes in related experiments (**Figure S1A**).

Consider a simple developmental process that occurred in a temporal interval of *d*, during which cells transitioned from state *i* to state *j* happened at a rate of *T_i_*_,*j*_(*d*) per unit time. The cell phylogeny of such a developmental process allows the phylogenetic adjacency between cells in state *i* and state *j* at a phylogenetic distance of *d*, or *A_i_*_,*j*_(*d*), to be directly calculated as the fraction of cell pairs at lineage distance *d* that are in states *i* and *j* (**Figure S1B**). Assuming invertibility, it can be shown that *T_i_*_,*j*_(*d*)^2*d*^ = *A_i_*_,*j*_(*d*)⁄*p_i_*, where *p_i_* is the fraction of cells in state *i* in the population of cells. Combining *T_i_*_,*j*_(*d*) for all the *N* cell states appearing at the tips of the lineage tree, we would have a *N*×*N* matrix *T*(*d*), which can be estimated from the corresponding *N*×*N* phylogenetic adjacency matrix of *A*(*d*) (**Figure S1C**).

Additionally, if the transition rates change as development proceeds, *T*(*d*) should also change for different *d*. Assuming similar *d* has similar *T*(*d*), we infer the optimal *T̂* for a range of *d_l_* < *d* ≤ *d_u_*(*d_l_* > 0) by minimizing the sum of the Euclidean distances between the *T̂* and every *T*(*d*). It should be noted that *d* could theoretically be a continuous variable indicating the temporal duration (chronological time, number of cell generations, etc) of the developmental process under investigation, but in practice it is discretized due to the limited temporal resolution of the experimentally determined cell phylogeny (see below for our estimation of “normalized depth” of an internal node/subtree). PASTRI is similar to the standard KCA^1,44^ but without assuming a constant *T*, and it allows user-defined constraints on *T* to incorporate prior knowledge about the developmental process such as irreversible transitions (**Figure S1D**). Algorithmic details of PASTRI can be found. PASTRI’s source code, and usage examples are available on its GitHub repository.

### Evaluation of PASTRI by simulated developmental cell phylogenies

We simulated sets of cell phylogenies with one or two rounds of cell division giving rise to 1,000 terminal cells, during which cell state transitions were governed by predefined transition rate matrix(es) consisting of five distinct states. In the simplest model, where only one round of division is simulated, we generated 500 mother cells whose states are randomly selected from the five possibilities. Each mother of state *i* produces two descendants whose states are probabilistically defined by *T_i_*_,*j*_. In the two-stage model, there are two predefined transition rate matrices, *Ta* and *Tb*. There were 250 grandmother cells created with random states. Each grandmother of state *i* produces 2 descendants (mother cells) whose states are probabilistically defined by *Ta_i_*_,*j*_, followed by another round of division for each mother cell similarly governed by *Tb*.

On the basis of one set of simulated phylogenies, we calculated the transition rates per division using PASTRI. The inferred rates were compared to the true (predefined) rates using the Euclidean distance (i.e. Frobenius norm. See **Figure 2B/E/H/J**). To compare this observed distance with its null expectation, we generated 1,000 sets of randomized phylogenies by permutating the terminal cells, applied PASTRI to each set, and calculated the Euclidean distance between the PASTRI results and the true transition rates. By comparing the observed distance with 1,000 distances derived from randomized phylogenies, a *P* value was calculated (**Figure 2C/F/I/K**).

To examine how the cell sampling density of the lineage tree affects PASTRI accuracy, we simulated ten phylogenies each with 50,000 terminal cells using a random bifurcation model. That is, starting from a “tree” with a single terminal cell, bifurcation (cell division) is applied to one randomly chosen terminal cell, and this process is repeated until there are 50,000 terminal cells. Then the normalized depth of internal cells is defined as follows:

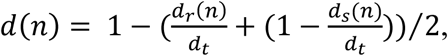

where *d_r_*(*n*) is the depth (number of internal nodes) from the root to node *n*, *d_s_*(*n*) is the maximum depth from node *n* to any of its descendant tips (i.e., the depth of the subtree rooted at *n*), and *d_t_* is the total depth of the phylogeny (i.e., the maximum root-to-tip depth). This definition normalizes all node depths to the interval of [0, 1], where 0 corresponds to the tips and 1 to the root. For the cell states on the tree, we assigned the root cell to state A and then simulate cell state transitions per cell division using a predefined transition matrix (**Figure S4A** left) for cells with normalized depth > 0.7 (the cells closer to the root), followed by another predefined transition matrix (**Figure S4A** right) for all remaining cells. As a result, we obtained a cell phylogeny with 50,000 terminal cells all annotated with cell states. From one original phylogeny of 50,000 terminal cells, we randomly selected 50%/20%/10% of the tips (corresponding to 25,000/10,00/5,000 cells) to construct a partial phylogeny. The cell states of the selected terminal cells were preserved in the partial phylogeny, but the normalized depths were recalculated using the partial phylogeny. We compared the PASTRI results derived from the full and partial phylogenies with the true (predefined) rates (rescaled to the time unit of *d*=1) using Euclidean distances (**Figure S4B**). Each level of subsampling was conducted with 100 random subsamples.

### Evaluating PASTRI with the developmental cell phylogenies of *C. elegans*

Fluorescence reporters have previously^20,21^ been used to determine cell lineage-specific expression patterns of 128 marker genes during the development of *C. elegans*. The data were downloaded along with the underlying cell phylogenies (lineage trees) from the EPIC database (Expression Patterns In Caenorhabditis)^20,21^. As the cell phylogenies of different genes were traced up to different end-points, we analyzed each gene individually using its corresponding cell phylogeny. Thirteen genes were removed at this step because their cell phylogenies were too small (< 200 terminal cells), giving rise to 115 genes analyzed below.

For simplicity, we ignored temporal variations in gene expression throughout a cell’s life cycle and used only the (chronologically) last measured value for the focal cell. A gene is considered active (ON) if its background-subtracted expression in a cell exceeds 30% of its maximal expression observed across the entire phylogeny, or inactive (OFF) otherwise. The normalized depths of internal cells are calculated as above-mentioned for the simulated trees, where all node depths are normalized to the interval of [0, 1], with 0 corresponds to the tips and 1 to the root.

To estimate the true transition rate, we first calculated the frequency of state changes along the targeted depth range. For example, for all branches spanning the depth range of 0.2-0.4, we determined the ratio between *N*_OFF_ON_, the number of branches that originate from any inactive mother cell and end at active daughter cell, and *N*_OFF_all_, the total number of branches that originate from any inactive mother cells. Summarizing the true frequencies of all kinds of state transitions as a matrix of *R*(Δ*d*), where Δ*d* = 0.4 − 0.2 = 0.2, we then scaled the true transition frequency to true transition rate by *R*(Δ*d*)^1/Δ*d*^, so that it became comparable with PASTRI inference in terms of time unit. Lastly, the true transition rates estimated from different depth intervals will be averaged into a final true transition rate.

PASTRI was used to estimate the transition rate from inactive to active state individually for each gene, using only the phylogeny and its state in terminal cells, but not that in internal cells. It is important to note that both PASTRI inference and the calculation of the true transition rate are not necessarily for the entire phylogeny, but can be limited to specific developmental stages (normalized depth in **Figure 3B-E**). To determine the accuracy of PASTRI inference at a given developmental stage, we calculated the Euclidean distance between the PASTRI-inferred and true OFF-to-ON transition rates of the 115 genes (**Figure 3C**). For the purpose of assessing statistical significance, 1,000 random phylogenies were generated by shuffling the terminal cells (the composition of cells in different states and the topology of the phylogeny remained unchanged). The accuracy of the 1,000 PASTRI inferences derived from these random phylogenies, i.e. their Euclidean distances to the true transition rates, were compared with that derived from the true phylogeny, by a permutation test (**Figure 3E**).

### Construction of the lineage tracer hESC cell line

To test PASTRI in a developmental process with prior knowledge, we used the substitution mutation-aided lineage-tracing (SMALT) system^24^ to reconstruct the cell phylogeny of an *in vitro* directed differentiation model from human embryonic stem cells (hESC) to hepatic stellate cells^22^. The SMALT system consists of an inducible mutator and a lineage barcode that is targeted by the mutator. The mutator is a doxycycline-inducible gene (hereinafter referred to as AI) that encodes a fusion protein comprising three components, namely AID, an activation-induced cytidine deaminase capable of creating C-to-T mutations on DNA, iSceI, a homing endonuclease that binds to an 18-bp DNA motif but with the endonuclease function inactivated, and UGI, a uracil-DNA glycosylase inhibitor counter-acting DNA-repair triggered by uracil N-glycosylation. The lineage barcode, which is the major readout for lineage tracing, is a 1376 bp segment of DNA containing a 10-bp integration barcode (intBC) and 16 iSceI binding motifs surrounded by AID-preferred sequences. Once integrated into the genome, the lineage barcode records the mitotic division history as heritable mutations induced by the mutator, allowing the recovery of the cell phylogeny via analyzing the mutated barcodes by substitution-based phylogenetic reconstruction algorithms.

A lineage tracer hESC cell line was constructed by genomic integration of the SMALT system, which was accomplished in three consecutive steps. The first step is the integration of AI. The doxycycline-inducible AI vector (PB-Tet-ON-T8>AI:T2A:mCherry-EF1A>Puro:T2A:rtTA, VectorBuilder:VB210226-1087xxf) was co-transfected with hyPBase (VectorBuilder:VB190515-1005nrp) in a ratio of 30 μg : 3 μg for 1×10^6^ cells by NeonTM transfection system (Life, MPK5000). To ensure adequate AI expression, Puromycin (1.0 μg/ml, InvivoGen, ant-pr-1) and Doxycycline (Dox, 1.0 μg/ml, Sigma, D9891) were applied simultaneously for 5 days, followed by two rounds of flow cytometry sorting for cells with strong mCherry signals.

The second step is the integration of lineage barcode. The lineage barcode vector (PB-CAG>eGFP:10bp-intBC:Barcode-EF1A>Bsd, VectorBuilder:Lib210303-1332qxe) was co-transfected with hyPBase in a ratio of 50 ng : 5 ng for 1×10^6^ cells by Neon^TM^ transfection system. After blasticidin (10 μg/ml, InvivoGen, ant-bl-1) selection, most transformants exhibited green fluorescence, supporting the transcriptional activity of the lineage barcode (**Figure S6A-B**). To verify that the lineage barcode was integrated as a single copy, transformant cells were individually inoculated onto 96-well plates and cultured for twelve days (1.0 ug/ml Dox treatment since day 2). Sixteen healthy colonies were selected and subjected to genomic DNA (gDNA) extraction, lineage barcode amplification with Phanta Max Super-Fidelity DNA Polymerase (Vazyme, P505) and primers T-Bar-F/T-Bar-R, TA-cloning with pCE-Zero (Vazyme, C115), colony PCR, and Sanger sequencing for 8 TA-clones. We recovered unique intBC from 14 out of the 16 colonies, suggesting that most cells have a single copy lineage barcode. Using primers were listed, the expression levels of AI, rtTA and lineage barcode transcripts were further confirmed via PCR followed by electrophoresis (**Figure S6C**), as well as RT-qPCR (**Figure S6D**).

The third step is optimizing the stability and efficiency of the SMALT system in hESC. We inoculated individual colonies of ∼ten cells into 96-well plates and cultured for ∼eight days. Each colony exhibiting normal morphology and uniformly strong green (eGFP) fluorescence was transferred to two replica wells on a 24-well plate. In one of the replicas, 1.0 µl/ml Dox was added to induce AI and mCherry expression for four to seven days. If both mCherry and eGFP fluorescence signals were bright and relatively uniform, the gDNA from all cells in that well was extracted and subjected to lineage barcode amplification, TA-cloning and Sanger sequencing as described above. The efficiency of editing in this well was then evaluated by the sequences of 24 TA-clones. Two wells with the highest editing efficiency were selected, and the cells in the other Dox-free replica of them were frozen and preserved for later use. Single-copy integration of the lineage barcode was confirmed by Sanger sequencing of the intBC (**Figure S6E**).

Additionally, we examined the editing efficiency of colonies seeded with a small number of initial cells. In four 96-well dishes, matrigel (Corning, 354277) was plated and each well was seeded with one lineage tracer hESC manually by micro-manipulation. The cells were cultured in 100 µl of mTesR media (Stemcell, 85850) for 12 days, to which 10 µl of CloneR (Stemcell, 05888) were added on day 0 and day 2, and 1.0 µg/ml Dox^+^ mTesR media was added and refreshed every 48 hours since day 2. Normally surviving colonies after the 12-day culture were harvested using Accutase (Stemcell, 07920). Next, genomic DNA was extracted from each colony using the QIAamp DNA Micro Kit (Qiagen, 56304) and PCR amplified for the lineage barcode. Single-copy integration of the lineage barcode is confirmed by Sanger sequencing of the intBC, and AI-induced mutations accumulated during colony formation were identified via TA-cloning followed by Sanger sequencing (**Figure S6F/G**).

### Directed differentiation from hESC to HSC

We adapted a previously described protocol^22^ for *in vitro* directed differentiation of hepatic stellate cells (HSCs) from hESCs. Briefly, 12-well plates were coated with a 1:50-reduced growth factor Matrigel (Corning, 356230) and low-glucose DMEM medium (GIBCO, 11885084) at 37°C for 30 minutes and washed with DPBS before use. At 70-90% confluency, hESC were passaged by incubating with Accutase cell dissociation reagent for 5 minutes at 37°C. Cell suspensions were diluted at a 1:2 ratio with DPBS, pipetted gently to mechanically break any clumps into single cells, and pelleted by centrifuging at 350 g for 3 minutes. The hESCs were plated in complete mTeSR medium supplemented with Y27632 (10 μM) (Stemcell, 72304) in 12-well plates at a density of ∼62,000 cells per square centimeter. The following day, the medium was changed to mTeSR without ROCK inhibitor. The differentiation was induced the next day with HSC differentiation medium consisting of 57% low-glucose DMEM medium, 40% MCDB-201-water (USBiological, C4000-05), 0.25x linoleic acid-bovine serum albumin (Sigma, L9530), 0.25x insulin-transferrin-selenium (GIBCO, 41400045), 1% penicillin streptomycin (LONZA, 17-603E), 10^-4^ M L-ascorbic acid (Sigma, A8960), 2.5 mM dexamethasone (Sigma, D1756) and 50 mM β-mercaptoethanol (GIBCO, 21985023). Growth factors were added as follows: BMP4 (R&D, 314-BP-010) (20 ng/ml) from day 0 to day 4, FGF1 (R&D, 232-FA-025) and FGF3 (R&D, 1206-F3-025) (20 ng/ml) from day 4 to day 8, and retinol (Sigma, R7632) (5 mM) and palmitic acid (Sigma, P0500) (100 mM) from day 6 to day 12 (**Figure S7A**).

To verify the success of directed differentiation, cells were dissociated into a single-cell suspension (FACS buffer, 2% FBS in DPBS) and stained at 4 °C for 30 minutes with antibodies for PDGFRβ (PE-CD140b clone 28D4, BD, Cat#558821, 1:100; PE mouse IgG2a, κ isotype, Biolegend, Cat#400213). Dead cells and debris were excluded based on scatter characteristics during flow cytometry analysis, and gating was determined by using isotype controls. Stained cells were analyzed with Attune NxT Flow Cytometer and official software (Thermo Fisher Scientific). Vitamin A-storing cells were similarly detected by flow cytometry using the autofluorescent at 350– 450/50-A, with the hESC cells as a negative control.

### The cell phylogeny of the hESC-to-HSC directed differentiation

We aimed to simultaneously evaluate single-cell transcriptomes and cell phylogenies for the directed differentiation from hESCs to HSCs, a stage at which there are <10,000 cells in the colony, so that a large proportion of cells can be sampled. To prepare the ancestor hESCs, colonies were cultured in mTesR media for 4-6 days and induced by Dox during the last two days, so that the hESC colonies were marked by primary editing events (to distinguish ancestor cells). After digestion by GCDR (stemcell, 07174), we resuspended them at a density of 1-2 colonies/μl. By micromanipulation, individual colonies (∼10 cells per well) were manually inoculated in 96-well dishes coated by Matrigel 1:50 diluted with low-glucose DMEM. Complete mTeSR medium supplemented with CloneR (10:1) was added in the first 48h to promote the survival of the cells. Directed differentiation was then initiated by applying both Dox (1.0 μg/ml, to induce the lineage barcode mutation) and the differentiation reagents described above to the surviving colonies, except that it was stopped on the 8^th^ day after initiation. Finally, colonies with intermediate size (∼3,000 cells) and >80% EGFP^+^mCherry^+^ cells were digested with 0.25% trypsin-EDTA (GIBCO, 25200056) for 3-5 minutes at 37°C, washed in DPBS containing 0.04% BSA, filtered with a cell strainer of 40 μm, centrifuged at 500 g for 5 min at 4 °C, and resuspended in single cell resuspension buffer (0.04% BSA in DPBS). For undifferentiated hESC samples, we first dissociated the normal hESC colonies into single cells using Accutase, counted the cells, and then resuspended them at a density of 1-2 cells/μl. Next, the single cells were directly plated into 96-well dishes and cultured in mTeSR medium for 12 days. During the first 4 days, CloneR was supplemented in the medium, and starting from the 2nd day, Dox (1.0 μg/ml) was added to induce lineage barcode mutations. Finally, the colonies were digested with Accutase at 37°C for 5 minutes, washed in DPBS containing 0.04% BSA, centrifuged at 500 g for 5 minutes at 4 °C, and resuspended in single cell resuspension buffer. The single cell suspensions were loaded into the 10x Genomics Chromium Single Cell chips, DNA libraries were prepared using the Chromium Single Cell 3’ GEM Library & Gel Bead Kit v3 (10x Genomics, PN-1000269) according to the manufacturer’s instructions. Each cDNA library was split into two halves, with the first half undergoing conventional RNA-seq by NovaSeq for single-cell transcriptomes, and the other half undergoing PCR amplification of the lineage barcode using KOD FX Neo (TOYOBO, KFX-201), followed by HiFi sequencing of the lineage barcode using PacBio Sequel II. The amplification of the lineage barcode was conducted with 50μl of solution with the following procedure: (i) pre-denaturation at 94°C for 2 minutes; (ii) denaturation at 98℃ for 10 seconds, annealing at 60℃ for 30 seconds, extension at 68°C for 1 minute, with these steps being cycled 25 times; (iii) incubation at 68°C for 10 min to ensure full extension, followed by holding at 4°C.

For the single-cell transcriptomes, raw sequencing reads were aligned to the human genome (GRCh38) and quantified using Cell Ranger^45^ (v7.1.0). Cells with mitochondrial content exceeding 25% and expressing fewer than 200 genes were excluded using Seurat^46^ (v4.4.0) in R (v4.2.3). Doublets were identified by DoubletFinder^47^ and removed. Consequently, 6,683 and 26,755 single-cell transcriptomes were obtained from HSC samples and hESC samples, respectively. To identify cell types/states by clustering, while avoiding biases due to unbalanced cell numbers from the differentiating and non-differentiating samples, we randomly selected the same number of cells from hESC samples as that from HSC samples. The balanced samples were subjected to Single-cell Orientation Tracing (SOT)^48^ to detect highly variable genes, and batch effect correction by Harmony^49^. Next, cells were clustered based on cell-cell distances estimated by FindNeighbors and FindClusters^46^. We obtained DEGs at various stages of differentiation towards HSCs from related single-cell^26^ (D4, D6, D8, D12 in **Figure 4E left**) and microarray datasets^22^ (D0 and D12 in **Figure 4E right**, bimod likelihood-ratio test, adjust *P* < 0.05). Cell clusters were scored based on the average expression and number of expressed stage-specific DEGs. Finally, we annotated the clusters based on both the inferred order of their appearance during differentiation and the presence of specific marker genes.

To reconstruct cell phylogeny based on the lineage barcode, raw subreads from PacBio Sequel II were aligned to the reference barcode using BLASR (release v5.1) and classified as positive or negative strand. Circular consensus sequences (CCS, also known as HiFi-sequencing reads) were generated for each strand in each Zero-mode waveguide (ZMW) by CCS (v6.4.0) with the parameters “-j=30 --min-passes=3”. Further confirmation of the identity of the CCS was achieved through the presence of three sub-sequences (“TGGACGAGCTGTACAAGTAA”, “AGATCGGAAGAGCGTCGTGTAG”, and “TCTCTATACGATCCGGACCT”) within it, allowing mismatches of up to two nucleotides. A 16-bp cell barcode, a 12-bp unique molecular identifier (UMI) and a lineage barcode were extracted from each CCS, which was then assigned to a specific cell if the UMI and the cell barcode from Sequel II and NovaSeq differed by no more than one nucleotide. A summary of the read numbers throughout the analyses above is presented. To further correct the sequencing errors, the lineage barcodes with the same cell barcode and the same UMI were merged by multiple sequence alignment via MUSCLE^50^ (v3.8.1551), followed by selecting the nucleotide with the highest frequency at each site. Such constructed lineage barcode transcripts were kept if captured by at least two ZMWs, and then subjected to mutation calling by comparison with the reference lineage barcode, allowing accumulated insertion and deletion up to 20 nucleotides in total. We then summarized the frequencies (number of unique UMIs) of different lineage barcode alleles from the same cell. To avoid the transcripts of the actual lineage barcode allele being overwhelmed by yet-to-decay transcripts of its ancestral alleles within the same cell, we identified all alleles whose substitutions were proper subsets of a focal allele, and added their frequencies to that of the focal allele. The lineage barcode allele of the focal cell was defined as the allele with the highest frequency, or the allele with a higher priority score (equal to the number of editing events multiplied by the number of ZMWs that capture it) in case of equal frequencies. Finally, we reconstructed the multifurcating cell phylogeny using the GTR+FO+I+R10 model in IQ-TREE, which is based on maximum likelihood^24^. We ran 1,000 rounds of ultrafast bootstrap approximation (-B 1000) and 1,000 rounds of SH-like approximate likelihood ratio test (-alrt 1000) to evaluate the quality of the consensus tree. A summary of the cell numbers throughout the analyses above was presented.

### The A549 cell culture and cell phylogeny

We also reconstructed the phylogeny of lung cancer cells using the SMALT system. By using the Neon™ transfection system, we integrated the AI vector and lineage barcode vector into the A549 cell line in a similar manner to the construction of the lineage tracer hESC cell line descrbied above. The resulting A549-SMALT cells were used to initiate single-cell seeded cultures in DMEM medium (GIBCO, C11330500BT) supplemented with 10% FBS (Corning, 35-081-CV) and Dox (1.0 µg/mL), which lasted for 14 days. For single-cell transcriptome analysis, we identified cell types/states using the default parameters in Seurat. During lineage tree reconstruction, we allowed up to 100 nucleotide insertions and deletions in total to accommodate the greater sequence difference observed among lineage barcode alleles captured for the A549 culture, which is presumably caused by a weaker DNA repair mechanism in cancerous cells. A summary of the cell numbers throughout the analyses was presented.

### PASTRI analyses on genomic barcoding-based cell phylogeny

PASTRI was applied to three genomic barcoding-based cell phylogeny datasets to estimate cell state transition rates, including (i) three hESC-to-HSC phylogenies derived from *in vitro* directed differentiation obtained in this study as described above, (ii) three hESC-to-PLP phylogenies reconstructed by a scGESTALT-like method previously described^35^, and (iii) two cell phylogenies determined for the clonal expansion of the human lung cancer cell line A549.

For cell phylogenies reconstructed by genomic barcoding-based methods, a proper metric for the depth of the MRCA of any tip pair is required for PASTRI inference. These phylogenies differ from the *C. elegans* cell phylogenies, in that the number of divisions along each root-to-tip lineage is highly heterogeneous, which may be the result of biological and/or technological stochasticity. To get a more robust depth estimation, we reasoned that the depth of an internal node (MRCA) can be reflected both by its distance from the root and its distance from the descendent tips, and therefore defined the normalized depth of a node as:

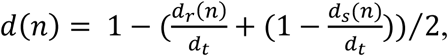

where *d_r_*(*n*) is the depth (number of cell divisions) from the root to node *n*, *d_s_*(*n*) is the maximum depth from node *n* to any of its descendant tips (i.e., the depth of the subtree rooted at *n*), and *d_t_* is the total depth of the phylogeny (i.e., the maximum root-to-tip distance). This definition normalizes all node depths to the interval of [0, 1], where 0 corresponds to the deepest tips and 1 to the root. This definition is also identical to that used for the downsampled simulated phylogeny (i.e., **Figure S4**).

Following the calculation of the normalized depths of internal nodes, each cell phylogeny was individually subjected to PASTRI, with prior knowledge^27,28,35^ of the developmental process incorporated as a constraint (irreversible differentiation of HSCs in the order of E-aHSCs, M-aHSCs, L-aHSCs and sHSCs; irreversible differentiation of PLP in the order of CD47^hi^, CD47^low^, GATA6^hi^SHH^hi^CD47^low^, GATA6^low^NKX2-1^neg^SHH^neg^CD47^neg^, GATA6^hi^NKX2-1^hi^CD47^hi^/GATA6^hi^). The reproducibility of PASTRI results is primarily measured by the Euclidean distance (Forbenius norm) between transition rate matrices inferred with phylogenies from different biological replicates. We also presented the between-replicate Pearson’s correlation coefficients of the inferred transition rates. Pearson’s correlation isn’t considered as the main measure of reproducibility since it is inflated by a few state pairs with high transition rates. To assess the statistical significance of the observed reproducibility, we extracted each pair of real phylogenies, say *A* and *B*, and then shuffled (within the phylogeny) the terminal nodes to create 1,000 randomized *A’* and 1,000 randomized *B’*. The composition of cells in different states and the topology of the phylogeny therefore remained unchanged. The real *A*-*B* reproducibility is compared with its null expectation assessed by 1,000 randomized values, each calculated by averaging one *A’*-*B* reproducibility and one *A*-*B’* reproducibility. *P* values from permutation tests are reported.

### PASTRI analyses on a lineage-traced lung adenocarcinoma dataset

To evaluate whether the cell-state transition rates estimated from low-sampling-rate datasets are sufficiently robust, we analyzed genetically engineered mouse model (GEMM) datasets of lung adenocarcinoma^40^. PASTRI analyzed phylogenetic trees and single-cell transcriptomes derived from two KP primary lung tumor samples (3435_NT_T3 and 3435_NT_T4), inferring a cell-state transition matrix for each sample. As comparisons, we also inferred the transition rates using KCA^44^, Fitch^15,16^, and PhyloVelo^17^, each run with the default parameters specified in the original publications.

## Data availability

The new data generated in this study were deposited to NCBI BioProjects under accession number PRJNA1424356 (hESC-HSC cell phylogeny), PRJNA1428415 (A549 cell phylogeny). Raw data underlying the hESC-PLP cell phylogeny were previously deposited to NCBI BioProjects under accession number PRJNA1099925. Data from Expression Pattern In Caenorhabditis (EPIC) is downloaded on May 1^st^, 2025 from https://epic.gs.washington.edu/.

## Code availability

All custom codes for processing the data are available at https://github.com/Yangwj01/PASTRI.

### Acknowledgments

We thank Jianzhi Zhang, Zheng Hu, Martin Tran, Michael B. Elowitz and Shou-Wen Wang for inspiring discussions and comments on the manuscript. This work was supported by the National Key R&D Program of China (grant numbers 2021YFA1302500 and 2021YFF1200904 to J.-R. Y., 2022YFA1106700 to F.C.), the National Natural Science Foundation of China (grant numbers 32122022 and 32361133555 to J.-R. Y.)

## Author Contributions

W.Y., Z.L., X.Y., P.W., X.Z. contributed equally to this work. J.Z., X.C. and J.-R.Y. conceived the idea, and designed and supervised the study. W.Y., P.W., C.R. conducted experiments and acquired data. K.L., J.C., F.C., X.H., J.Z., X.C. and J.-R.Y. contributed new devices/reagents/analytic tools. W.Y., Z.L., X.Y., P.W., X.Z., J.C., J.Z. and J.-R.Y. analyzed the data. W.Y., X.Y. and J.-R.Y. wrote the paper with inputs from all the authors.

## Supplemental Figures

**Figure S1.**
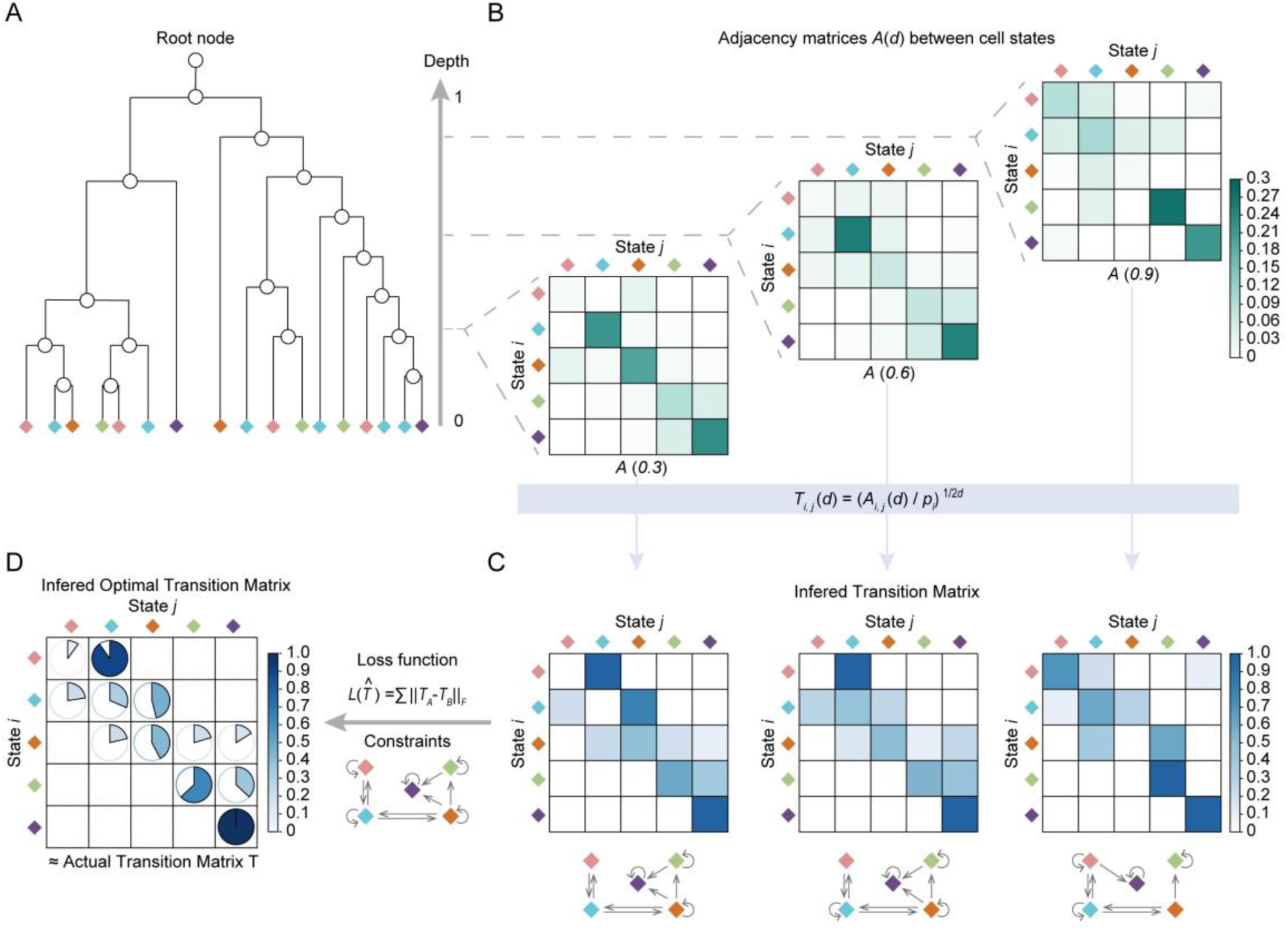
Schematic for the inference procedure of PASTRI. **(A)** PASTRI takes as input a cell phylogeny with terminal nodes (diamonds) annotated with cell states (colors). The divergence time between any two terminal nodes approximates to the depth of their most recent common ancestor (MRCA; internal node; circle). The depth is normalized to 0 (terminal) – 1 (root) (right axis). **(B)** PASTRI constructs an adjacency matrix *A*(*d*) at each depth by counting cell state pairs (states *i* and *j*) whose MRCA occurs at that depth. The color scale bar to the right indicates frequency. **(C)** From each adjacency matrix, PASTRI derives a matrix of transition rates per unit depth. **(D)** PASTRI finds the optimal transition matrix *T* by minimizing a loss function, within the predefined rate constraints. The loss function summarizes the differences between the candidate matrix and the depth-specific transition matrices for a chosen depth range.

**Figure S2.**
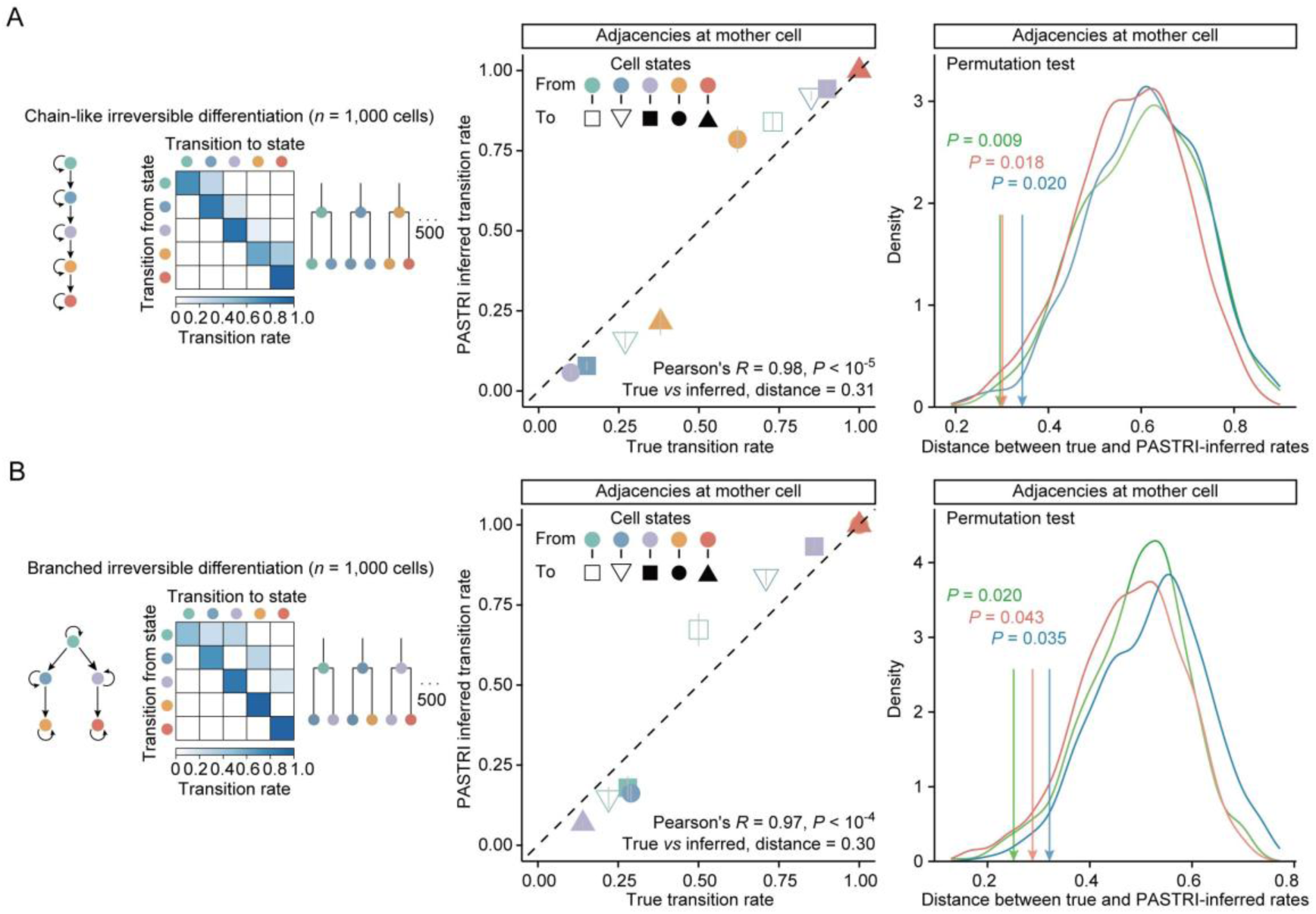
Results of testing PASTRI with cell phylogenies simulated with stable-irreversible dynamics. **(A)** A stable-irreversible chain-like transition model, depicted as a network and matrix in the left panel, was used to simulate 500 phylogenies of 2 terminal cells each (see Figure 2A). The middle panel illustrates a typical result from one simulation, comparing the true (*x*) and PASTRI-inferred (*y*) transition rates from state *i* (point color) to state *j* (point symbol). Bottom right corner shows the Pearson’s correlation test result and the Euclidean distance between the true and inferred transition rates (see Figure 2B). In the right panel, a colored arrow marks the observed Euclidean distance between the true and PASTRI-inferred transition rates. A curve of the same color shows the expected probability density of the Euclidean distance over 1,000 sets of phylogenies (500 per set) obtained by shuffling the terminal cell states. Three pairs of observed and expected Euclidean distances, from three independent simulation rounds, are shown in three colors (see Figure 2C). **(B)** Same as A, except for a stable-irreversible branched transition model.

**Figure S3.**
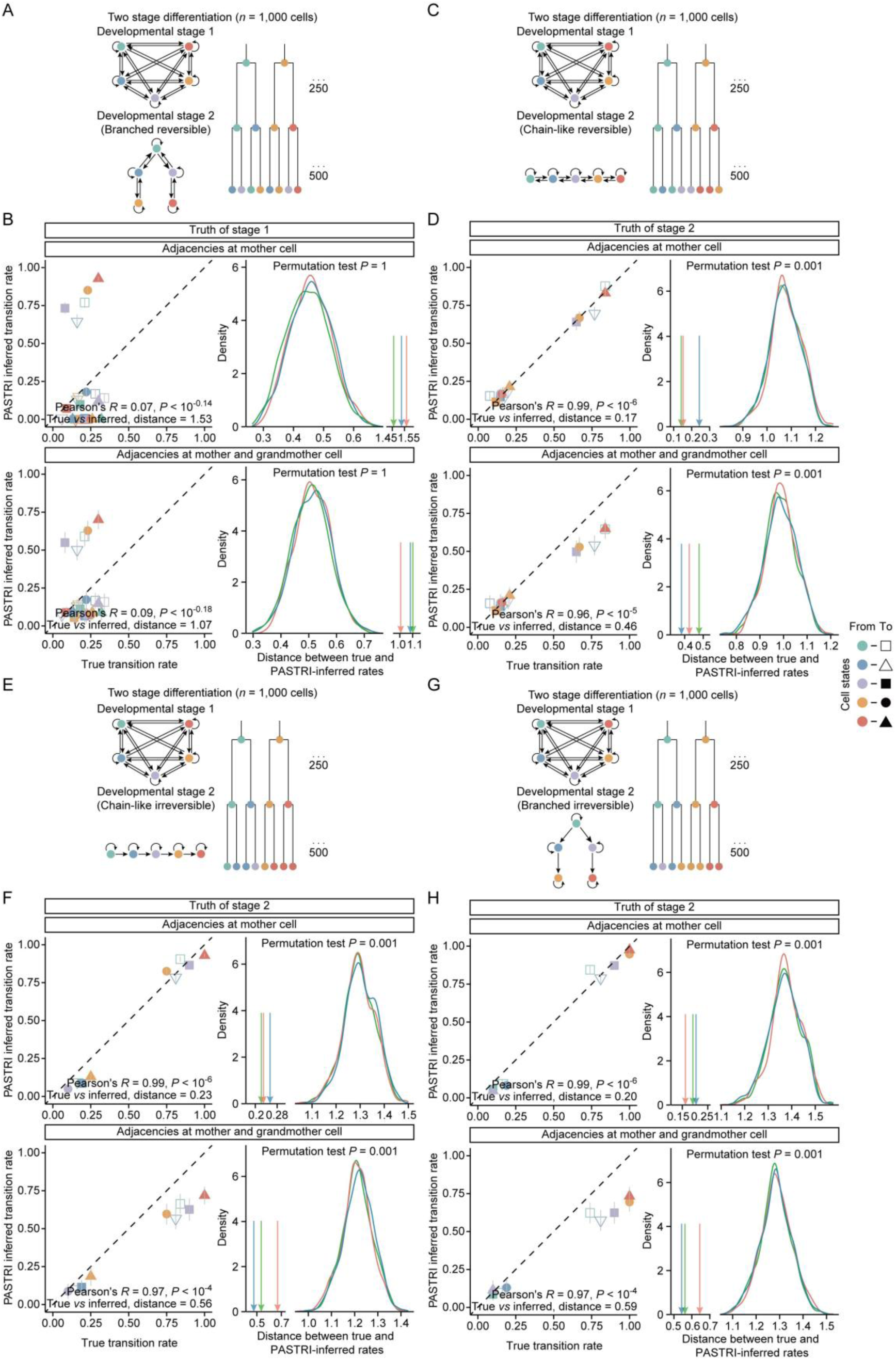
More results of testing PASTRI with simulated cell phylogenies. **(A)** Same as Figure 2H-I, except that PASTRI used adjacencies at both mother and grandmother cells, and was compared to the stage-1 truth. **(B)** Same as A, except that PASTRI was compared to the stage-2 truth. **(C)** A two-stage transition model with a chain-like reversible stage 2 was used to simulate 250 phylogenies each with two rounds of cell division, resulting in 1,000 terminal cells. **(D)** Same as Figure 2H-K, except that they are based on the transition model in panel **C**. **(E-F)** Same as panel **C-D**, except that they are based on a two-stage transition model with a chain-like irreversible stage 2. **(G-H)** Same as panel **C-D**, except that they are based on a two-stage transition model with a branched irreversible stage 2.

**Figure S4.**
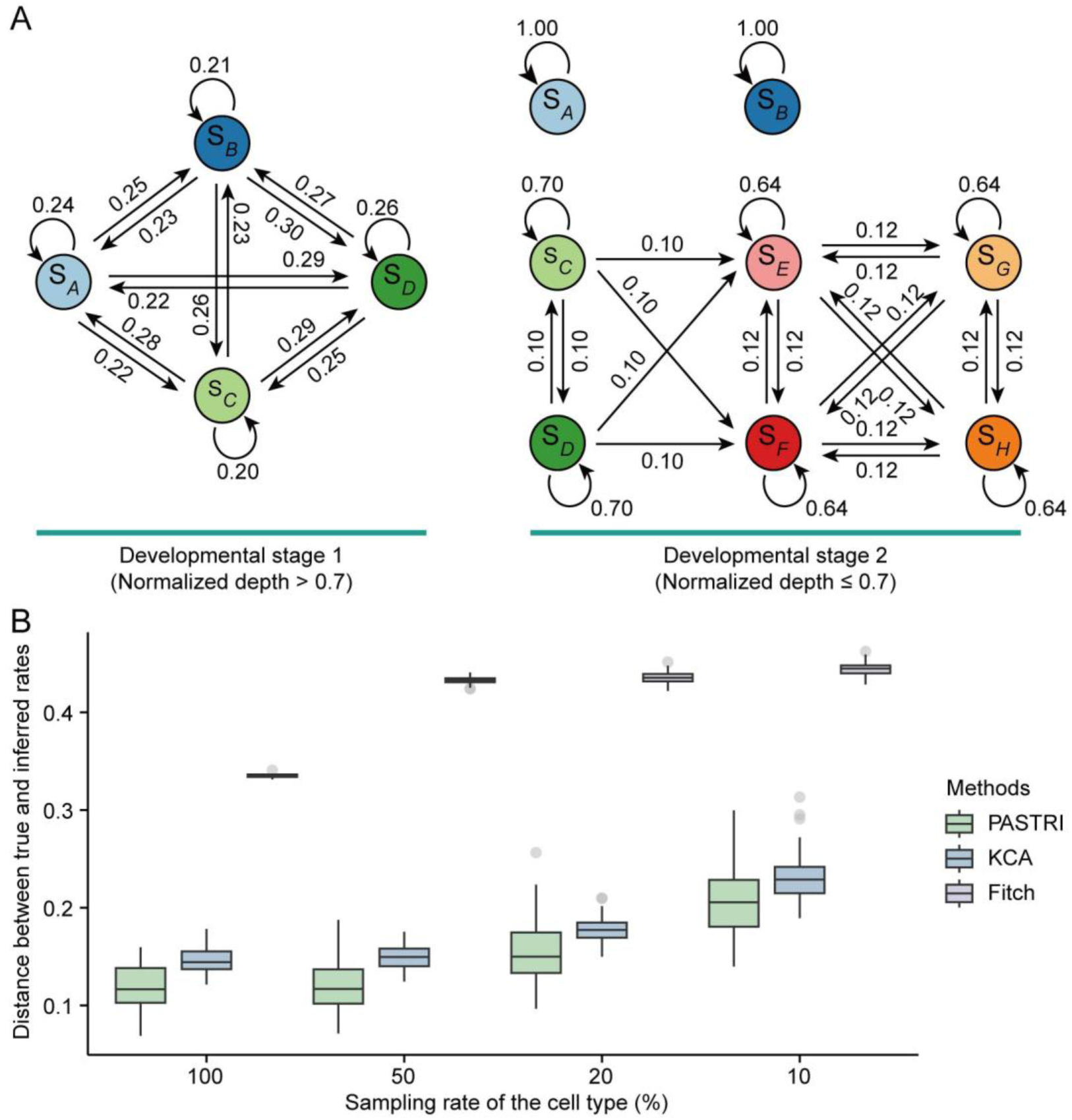
PASTRI outperforms other algorithms assuming stable transition rates regardless cell subsampling. **(A)** A two-stage transition model simulated one large phylogeny of 50,000 terminal cells, with the model switch occuring at *d* = 0.7 (See **Methods**). **(B)** Using the transition model in A, the Euclidean distance between the true (stage 2) and inferred transition rates (*y*) is compared across PASTRI (red), KCA (green), and Fitch (blue), with PASTRI remaining the most accurate despite cell subsampling (*x*) of the large phylogeny. PASTRI inferred most accurately at *d* ≤ 0.8 with 100% cell sampling, *d* ≤ 0.7 with 50% and 20% cell sampling, and *d* ≤ 0.5 with 10% cell sampling. The decreasing depth threshold reflects the depth underestimation expected by cell subsampling

**Figure S5.**
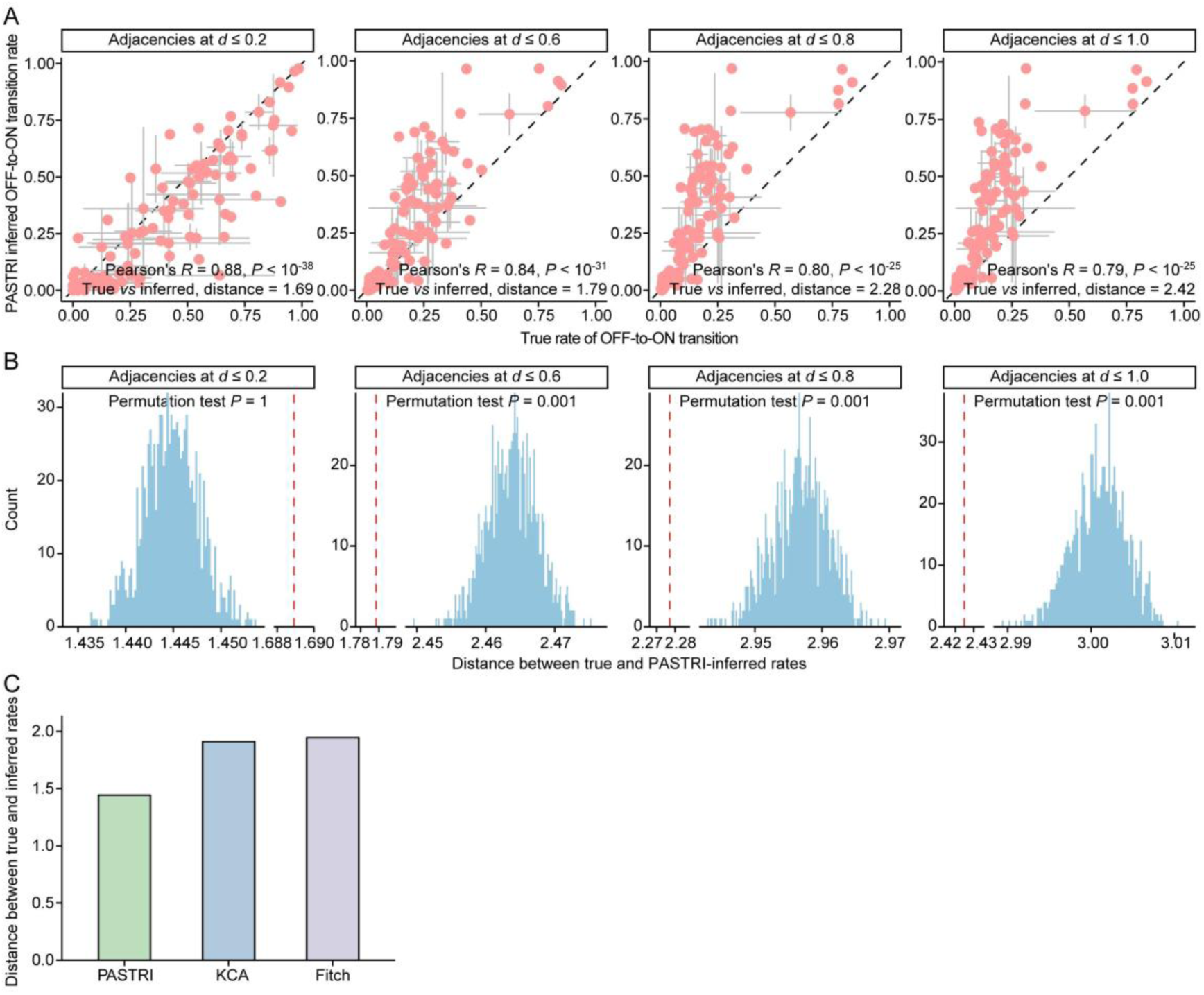
More results of testing PASTRI with cell phylogenies of *C. elegans*. **(A)** PASTRI-inferred OFF-to-ON transition rates (*y*) are compared with the true transition rates (*x*) across adjacencies up to different depth *d* (see **Methods**). Each panel shows the result for one depth value. Gray lines represent the standard error between biological replicates. Pearson’s correlation coefficient, *P* value, and Euclidean distance between the inferred and the true rates appear at the bottom of each panel. **(B)** The red dashed line marks the observed Euclidean distance between the inferred and the true rates. The blue histogram shows the expected probability density of the Euclidean distance over 1,000 randomized phylogenies obtained by shuffling the terminal cell states. *P* values from permutation tests are indicated on top. **(C)** Accuracy comparison of transition rate inference by PASTRI, KCA, and Fitch, inversely gauged by Euclidean distance between true and inferred transition rates.

**Figure S6.**
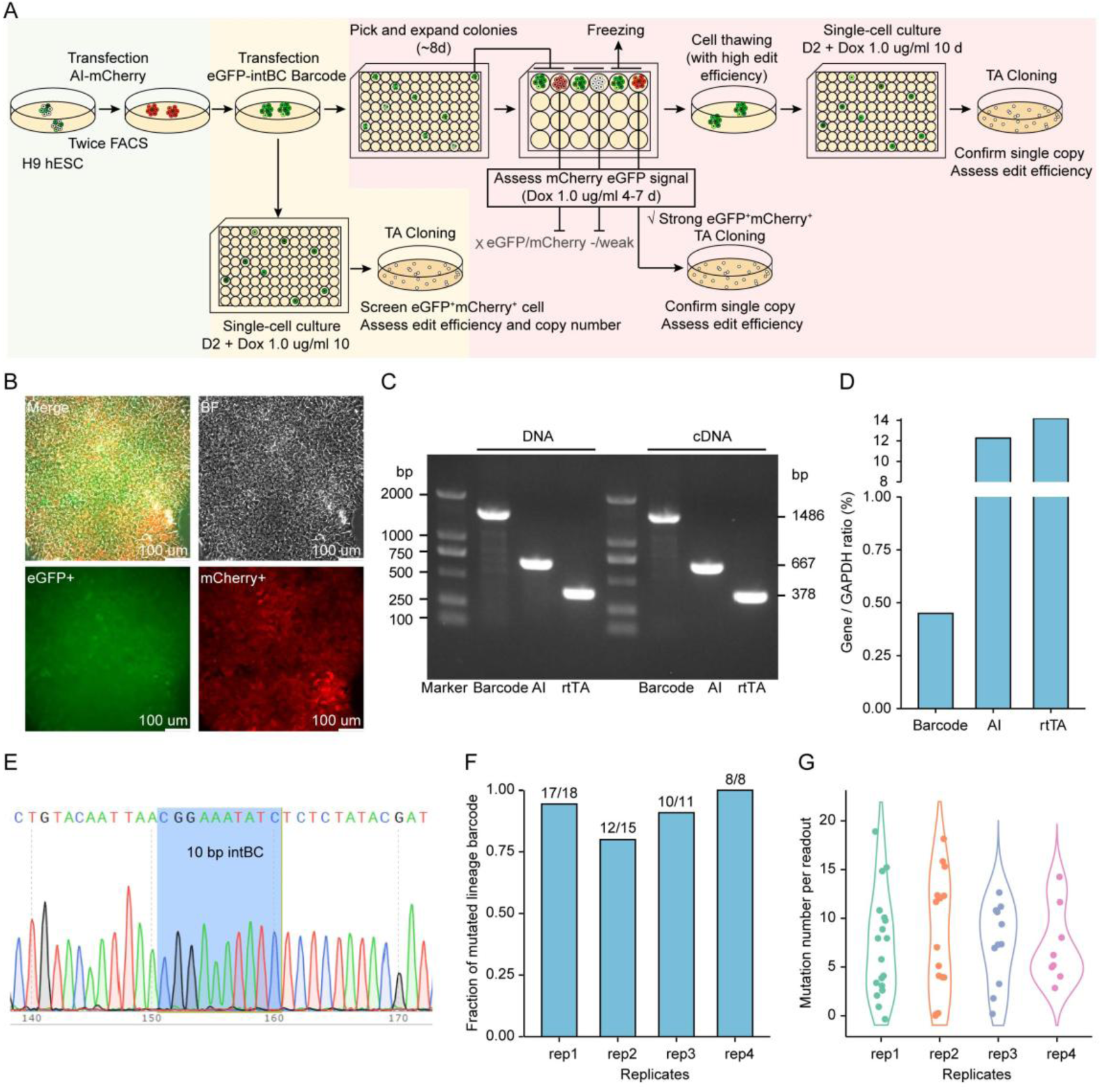
Construction of the SMALT lineage tracer hESC cell line. **(A)** Three-step construction workflow. **(B)** Typical colonies expressing lineage barcode (eGFP) and AI (SMALT mutator fused to mCherry). **(C-D)** Expression of lineage barcode and AI were confirmed via PCR followed by electrophoresis **(C)** and RT-qPCR **(D)**. **(E)** Typical sequencing results for the intBC, showing single copy integration of lineage barcode. **(F)** Fraction of mutated lineage barcodes, detected by TA clone. **(G)** Number of mutations per mutated lineage barcodes.

**Figure S7.**
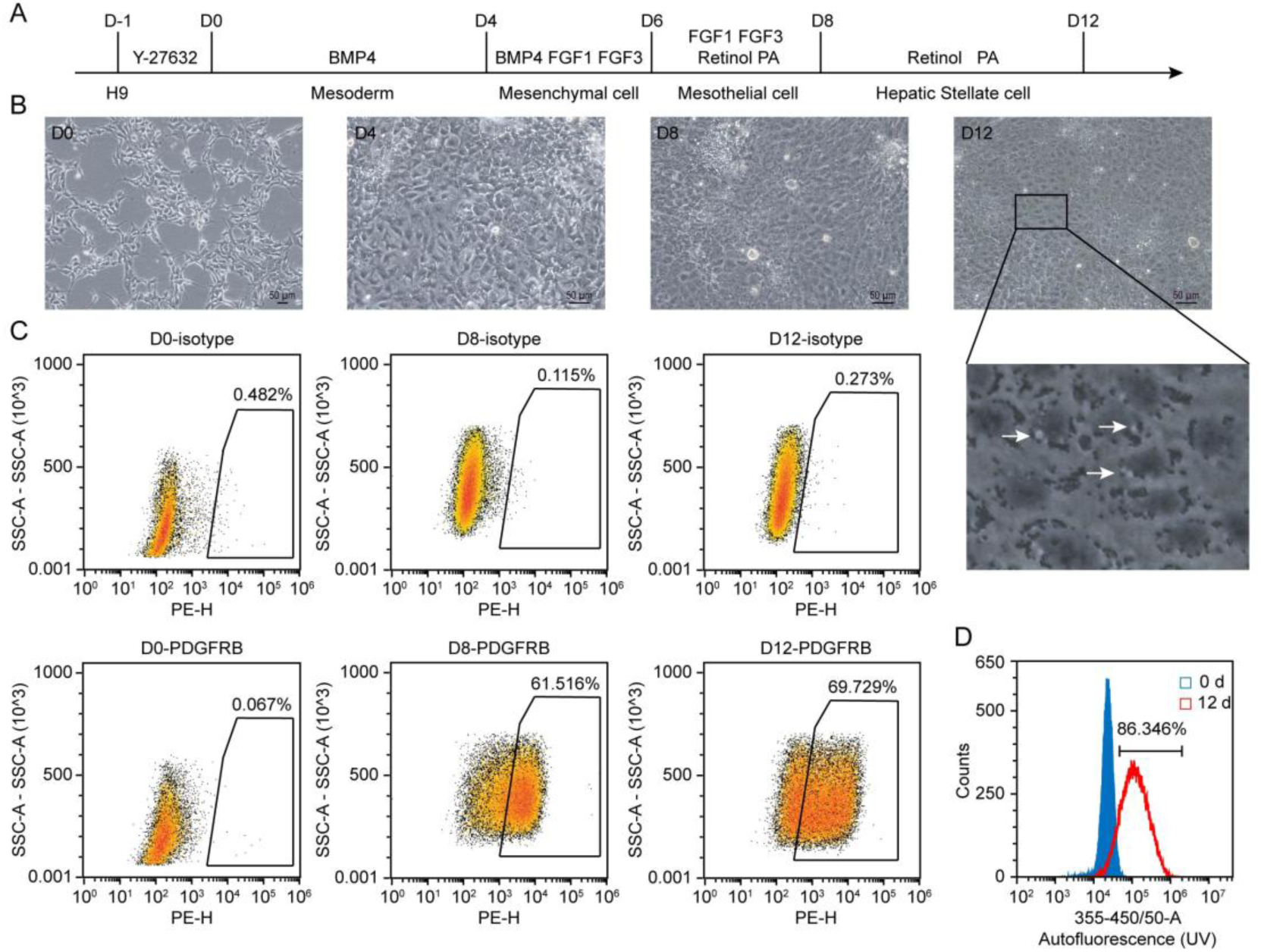
Directed differentiation from hESC to HSC. **(A)** Timeline of key treatments and cell state hallmarks during *in vitro* hESC-to-HSC differentiation. **(B)** Representative images of cell morphology on days 0, 4, 8, and 12 of differentiation. Zoomed-in view shows lipid droplets (white arrows). **(C)** Representative flow cytometry plots of PDGFRβ-positive cells on days 0, 8 and 12 of the differentiation. **(D)** Representative flow cytometry histogram of vitamin A expression at days 0 and 12.

**Figure S8.**
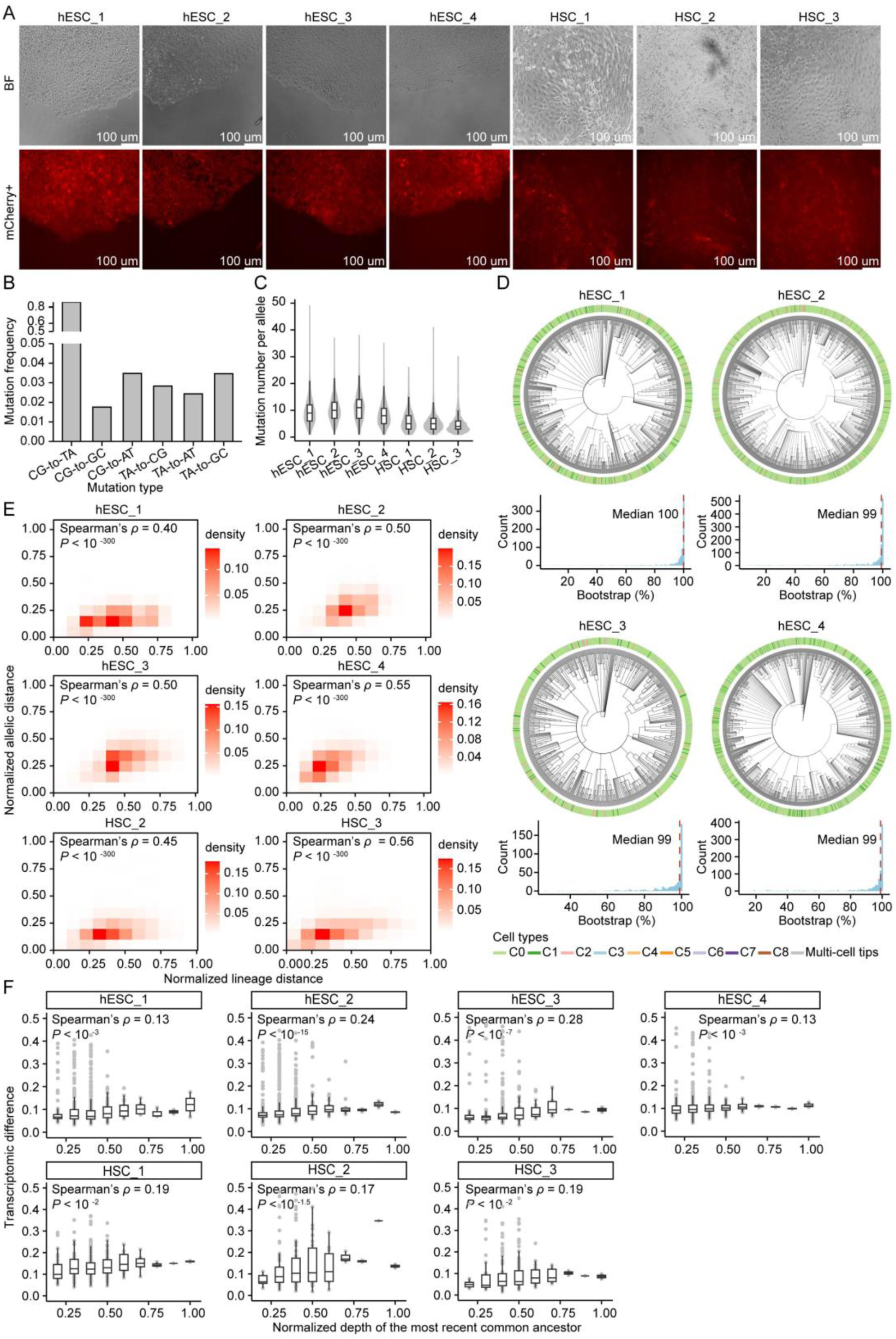
Quality of the SMALT lineage tracing. **(A)** Morphological and fluorescence imaging of hESC-to-HSC differentiation and hESC self-renewal cultures. **(B)** Mutation spectrum of the lineage barcode. **(C)** Per-allele mutation number for each sample, shown as violin plots with embedded box plots (center: median; hinges: first and third quartile). **(D)** Reconstructed cell phylogenies of hESC samples, with cell states of the terminal nodes colored on the outer ring. Inset histograms show the bootstrap support distribution across each phylogeny’s internal nodes. **(E)** Replicates Figure 4H for each sample (sample identity labeled above). **(F)** Cells closer in phylogeny share more similar transcriptomes (one sample per panel). For each subtree, transcriptomic differences (Pearson distances over genes expressed in > 50% of cells) are computed for all single-cell-tip pairs whose MRCA is the root of the focal subtree, then averaged. Box plots show the distribution of transcriptomic divergence across subtrees at each normalized depth (precision 0.1), with Spearman’s correlation coefficient and *P* value presented.

**Figure S9.**
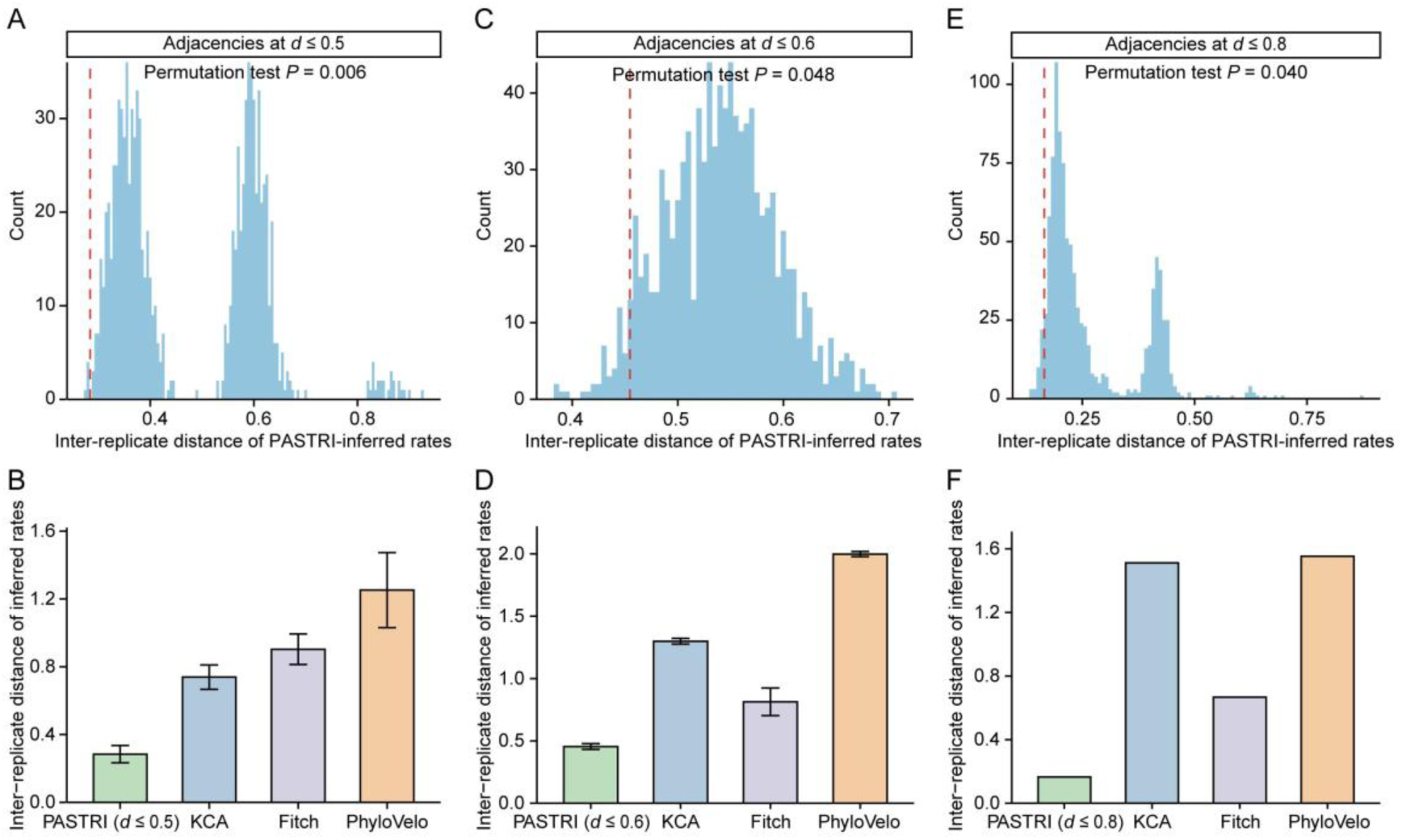
Precision of PASTRI inference across two lineage tracing systems and developmental *versus* cancerous cell phylogenies. (**A**, **C**, **E**) Permutation test of the Euclidean distance between PASTRI-inferred transition rates from biological replicates at specific adjacency depths (**A**: hESC-HSC, *d* ≤ 0.5; **C**: hESC-PLP, *d* ≤ 0.6; **E**: A549, *d* ≤ 0.8). Smaller distances (*x* axis) indicate more accurate inference. Red dashed line: PASTRI’s observed distance; blue histogram: null distribution from 1,000 randomized control phylogenies (shuffled terminal cell labels). (**B**, **D**, **F**) Accuracy comparison of transition rate inference by PASTRI, KCA, Fitch, and PhyloVelo, inversely gauged by Euclidean distance between transition rates inferred from biological replicates. Error bars represent standard error across replicate pairs.

**Figure S10.**
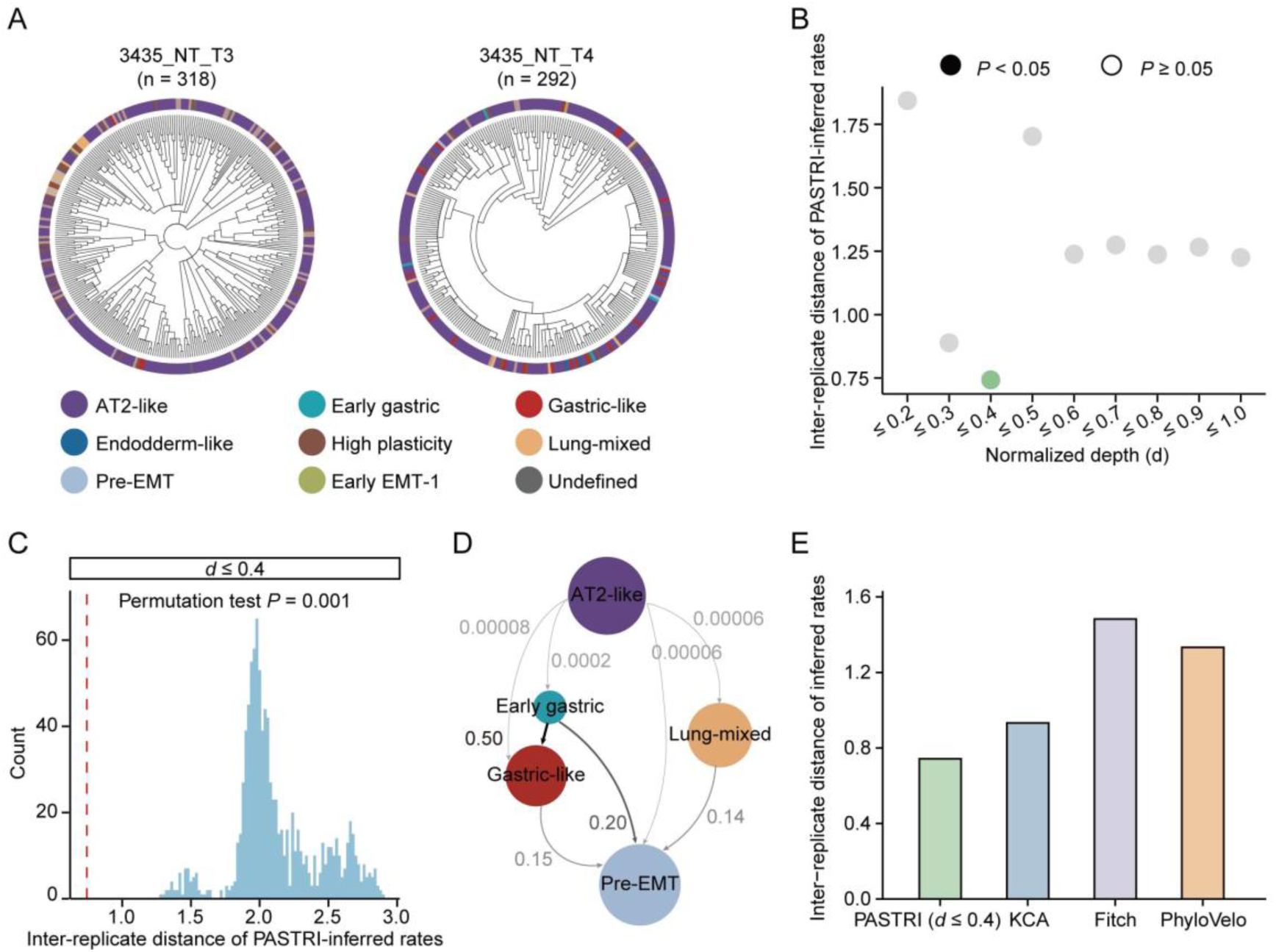
PASTRI captures cell fate transition dynamics for *in vivo* lung tumor evolution from mouse. **(A)** Phylogenetic trees from the KP mouse primary lung tumor, 3435_NT_T3 and 3435_NT_T4, in the Yang et al. dataset^40^. The scRNA-seq data, cell type annotations and lineage trees were obtained from the original study. **(B)** Among-replicate reproducibility (*y* axis) of PASTRI inferences versus adjacency depth *d* (*x* axis). Solid and hollow circles mark permutation *P* values < 0.05 and ≥ 0.05, respectively. **(C)** Permutation test of the Euclidean distance between PASTRI-inferred transition rates from biological replicates at specific adjacency depths (*d* ≤ 0.4). Smaller distances (*x* axis) indicate more accurate inference. Red dashed line: PASTRI’s observed distance; blue histogram: null distribution from 1,000 randomized control phylogenies (shuffled terminal cell labels). **(D)** Cell state transition graph averaged from the two biological replicates, inferred by PASTRI at *d* ≤ 0.4. **(E)** Accuracy comparison of transition rate inference by PASTRI, KCA, Fitch, and PhyloVelo, inversely gauged by Euclidean distance between transition rates inferred from biological replicates.

